# Endocrine-exocrine signaling drives obesity-associated pancreatic ductal adenocarcinoma

**DOI:** 10.1101/663583

**Authors:** Katherine Minjee Chung, Jaffarguriqbal Singh, Lauren Lawres, Kimberly Judith Dorans, Cathy Garcia, Daniel B. Burkhardt, Rebecca Robbins, Arjun Bhutkar, Rebecca Cardone, Xiaojian Zhao, Ana Babic, Sara A. Vayrynen, Andressa Dias Costa, Jonathan A. Nowak, Daniel T. Chang, Richard F. Dunne, Aram F. Hezel, Albert C. Koong, Joshua J. Wilhelm, Melena D. Bellin, Vibe Nylander, Anna L. Gloyn, Mark I. McCarthy, Richard G. Kibbey, Smita Krishnaswamy, Brian M. Wolpin, Tyler Jacks, Charles S. Fuchs, Mandar Deepak Muzumdar

## Abstract

Obesity is a major modifiable risk factor for pancreatic ductal adenocarcinoma (PDAC), yet how and when obesity contributes to PDAC progression is not well understood. Leveraging an autochthonous mouse model, we demonstrate a causal and reversible role for obesity in early PDAC progression, showing that obesity markedly enhances tumorigenesis, while genetic or dietary induction of weight loss intercepts cancer development. Bulk and single cell molecular analyses of human and murine samples define microenvironmental consequences of obesity that promote tumor development rather than new driver gene mutations. We observe increased inflammation and fibrosis and also provide evidence for significant pancreatic islet cell adaptation in obesity-associated tumors. Specifically, we identify aberrant islet beta cell expression of the peptide hormone cholecystokinin (CCK) in tumors as an adaptive response to obesity. Furthermore, beta cell CCK expression promotes oncogenic *Kras*-driven pancreatic ductal tumorigenesis. Our studies argue that PDAC progression is driven by local obesity-associated changes in the tumor microenvironment – rather than systemic effects – and implicate endocrine-exocrine signaling beyond insulin in PDAC development. Furthermore, our demonstration that these obesity-associated adaptations are reversible supports the use of anti-obesity strategies to intercept PDAC early during progression.

## INTRODUCTION

Pancreatic ductal adenocarcinoma (PDAC) is the third leading cause of cancer death in the United States and is expected to become the second within the next few years (Rahib et al., 2014; Siegel et al., 2019). Despite the development of combination chemotherapy, novel targeted small molecules, and immunotherapies that have revolutionized the care of many other cancers, long-term survival in PDAC remains low at <10% (Ryan et al., 2014; Siegel et al., 2019). In search of drug targets, genomic studies have identified frequent mutations in the proto-oncogene *KRAS*, occurring in >90% of human PDAC tumors (Bailey et al., 2016; The Cancer Genome Atlas Research Network, 2017). Unfortunately, no effective direct KRAS inhibitors are currently approved for clinical use (Papke and Der, 2017). Moreover, our recent work has demonstrated that PDAC cells can survive genetic ablation of *KRAS* both *in vitro* and *in vivo* (Chen et al., 2018; Muzumdar et al., 2017), arguing that resistance will subvert even the very best KRAS inhibitors. Alternative paradigms beyond genetic factors therefore need to be explored to develop novel therapeutic and preventative strategies in PDAC.

Epidemiologic studies in prospective cohorts have identified non-genetic host factors that contribute to PDAC development. In particular, body-mass index (BMI), a surrogate measure of obesity, has been shown to be positively correlated not only with an increased risk of developing PDAC (Larsson et al., 2007) but also more advanced disease at diagnosis and worse survival (Yuan et al., 2013). How obesity contributes to PDAC development and progression remains poorly understood. Previous research demonstrated that high-fat diet (HFD)-induced obesity can promote tumor growth or cancer progression in transplant or autochthonous models of PDAC (Incio et al., 2016; Khasawneh et al., 2009; Philip et al., 2013; Rooman et al., 2017; Zaytouni et al., 2017; Zyromski et al., 2009), and implicated dysregulation of fatty acid or nitrogen metabolism (Khasawneh et al., 2009; Zaytouni et al., 2017), COX2 activation (Philip et al., 2013), and aberrations in the desmoplastic response (Incio et al., 2016; Khasawneh et al., 2009; Rooman et al., 2017) as potential mechanisms. These studies were limited by the slow onset of obesity and lack of rapid reversibility in HFD models, as well as variations in fat content, types of fat, and age of administration in HFD protocols, all of which can confound results (Speakman, 2019). Importantly, these approaches provided no information on whether – and at what point – weight loss or other anti-obesity interventions might influence PDAC progression, despite significant implications for prevention and treatment (Massetti et al., 2017).

For this study, we develop a rapid and reversible autochthonous model of obesity-associated PDAC to overcome these important limitations. We demonstrate that obesity plays a stage-specific causal – and reversible – role in early PDAC progression. Molecular and histologic tumor analyses revealed prominent microenvironmental alterations, including increased inflammation and fibrosis plus evidence for marked pancreatic islet cell adaptation in the setting of obesity. We also identify aberrant expression of the peptide hormone cholecystokinin (CCK) in islet beta cells in tumors as an adaptive response to obesity and show that beta cell CCK overexpression itself promotes pancreatic ductal tumorigenesis. Together, these studies link obesity, changes in the local microenvironment, and tumorigenesis through a previously unappreciated endocrine-exocrine signaling axis in PDAC development.

## RESULTS

### Obesity drives pancreatic but not lung tumorigenesis

To model obesity-associated pancreatic cancer, we crossed leptin-deficient (*ob/ob*) mice with *KC* (*Pdx1-Cre; Kras^LSL-G12D/WT^*) mice predisposed to the development of precursor pancreatic intraepithelial neoplasia (PanINs) (Hingorani et al., 2003) to yield *KC; ob/ob* (*KCO*) mice (**Figure 1A**). Unlike HFD models, *ob/ob* mice exhibit early onset obesity due to impaired appetite suppression and decreased metabolism (Friedman, 2019). *KCO* mice were obese and developed increased primary ductal tumor burden (**Figures 1B-D**). They also showed enhanced progression to adenocarcinoma and markedly shortened survival compared to non-obese *KC* mice (**Figures 1E-F**). We similarly observed decreased survival using a distinct obesity model induced by loss-of-function mutations in the leptin receptor (*db/db*) (**Figures S1A-S1B**). These phenotypes were significantly more severe than seen in HFD models (Khasawneh et al., 2009; Philip et al., 2013; Rooman et al., 2017), corresponding to the degree of obesity. Consistent with human epidemiologic studies that demonstrate an association between obesity and pancreatic cancer, but not lung cancer (Lauby-Secretan et al., 2016), obesity induced by neither HFD nor *ob/ob* genotype shortened survival in lung cancer initiated by the spontaneous activation of a “hit-and-run” allele of oncogenic *Kras* (*Kras^LA2^*) (Johnson et al., 2001) (**Figures S1C-F**). Thus, these results support a specific effect of obesity on *Kras*-driven PDAC progression.

**Figure 1.**
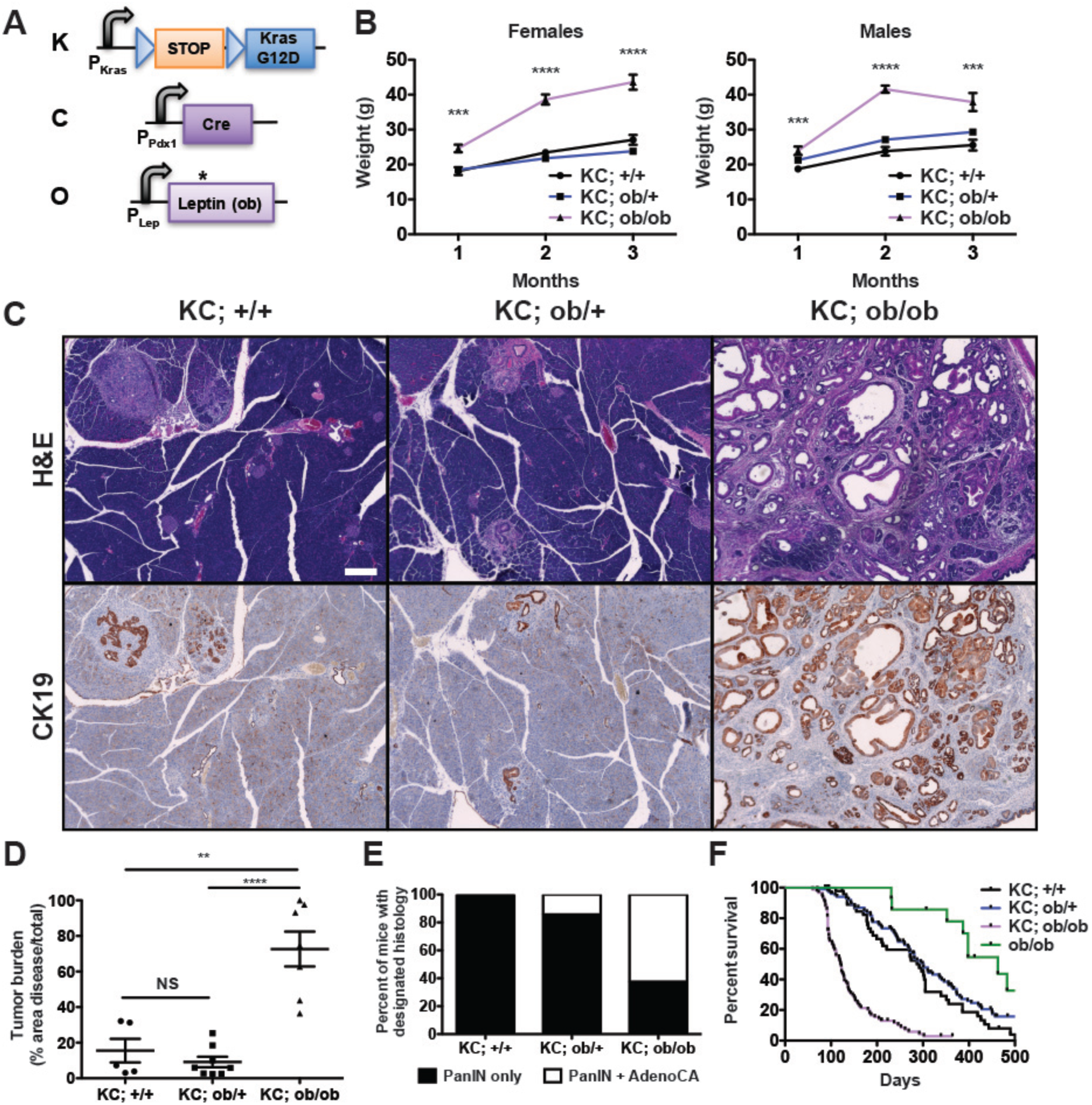
Accelerated PDAC progression in *KCO* mice. **A)** Schematic of transgenic/knock-in alleles in leptin-deficient *KC* mice (*KC; ob/ob* or *KCO*). Black arrows denote promoters. *Kras* and *Lep* promoters are endogenous. Blue triangles denote LoxP sites. A STOP cassette prevents oncogenic *Kras* (*G12D*) expression prior to Cre-mediated recombination. * denotes point mutation in *ob* gene causing premature stop. **B)** Average body weight +/− s.e.m. (in grams) of male and female *KC* mice of varying *ob* genotype over time (n=22-46 mice per group). ***p<0.001, ****p<0.0001 compared to *KC; +/+* mice, two-tailed student’s t-test at each time point. **C)** Representative histologic sections of pancreata from mice of designated genotypes at 3 months of age demonstrated differences in the generation of CK19-positive ductal tumors. Scale bar: 200 µm. **D)** Tumor burden (average % cross-sectional area of disease/total +/− s.e.m.) in mice of designated genotypes at 3 months of age (n=5-8 mice per group). **p<0.01, ****p<0.0001, two-tailed student’s t-test. NS = non-significant. **E)** Percentage of mice of particular genotypes harboring PanINs and/or adenocarcinoma by histology at 3 months of age (n=5-8 mice/group). **F)** Kaplan-Meier survival curves for non-obese *KC* mice (*KC; +/+* (n=51), non-obese *KC; ob/+* (n=106)), obese *KC* mice (*KC; ob/ob*, *KCO*, n=91), and obese non-tumor bearing controls (*ob/ob* (n=14)). Log-rank test: p<0.0001 *KC; +/+* vs. *KC; ob/ob*, p<0.0001 *KC; ob/+* vs. *KC; ob/ob*, p>0.05 *KC; +/+* vs. *KC; ob/+*.

### Weight loss intercepts early tumor progression in *KCO* mice

We chose the *ob/ob* obesity model because of the capacity for rapid reversal of the obesity phenotype through leptin restoration to study the effects of weight loss on PDAC development. We generated adeno-associated viruses (AAVs) that direct sustained leptin secretion (AAV-Leptin) or expression of green fluorescent protein (AAV-GFP) following a single intramuscular injection (**Figures 2A, S2A**, and **S2B**). AAV-Leptin reversed multiple phenotypes of leptin deficiency including obesity, hyperglycemia, and infertility (**Figures 2B** and **S2C-D**), as seen with leptin protein replacement (Friedman, 2019). Remarkably, early AAV-Leptin administration (following obesity onset but prior to significant tumor development) also impeded tumor progression proportional to the degree of weight loss (**Figures 2B-D**). In contrast, late AAV-Leptin administration (following advanced tumor development) induced weight loss but did not impact survival (**Figures S2E-F**). Moreover, recombinant leptin treatment of advanced murine PDAC cells *in vitro* induced on-target downstream signaling but did not alter cell viability (**Figures S2G-H**). These data support a stage-specific effect of obesity on early pancreatic cancer development. Experiments with caloric restriction argue that it is weight loss itself that impacts early tumor progression – rather than leptin signaling. Young *KCO* mice subject to caloric restriction exhibited significantly reduced tumor burden compared to *ad libitum* fed mice (**Figures 2E-F**), demonstrating the potential to halt or delay obesity-driven PDAC progression with weight loss.

**Figure 2.**
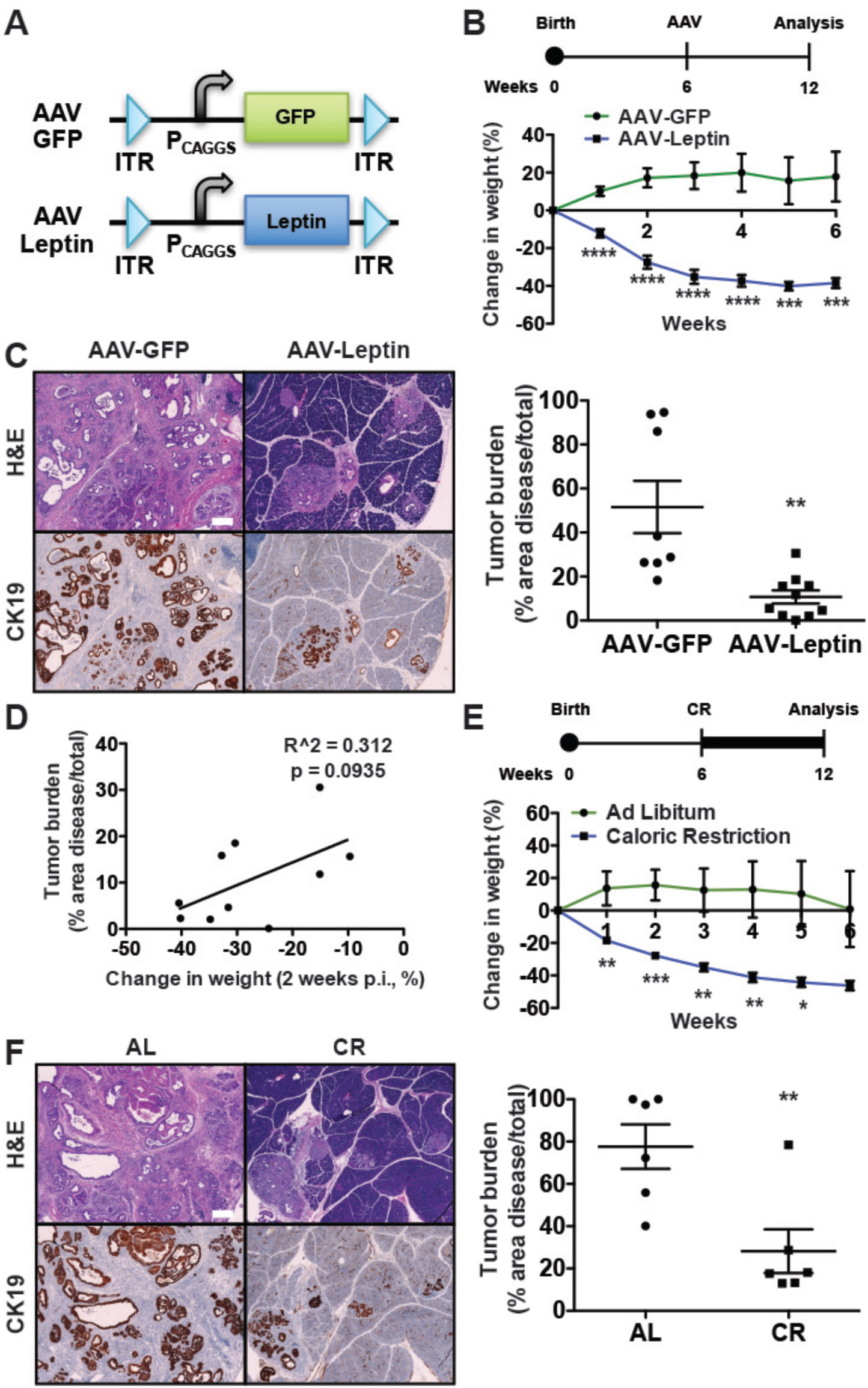
Weight loss intercepts early tumor progression in *KCO* mice. A) Schematic of AAV vectors administered to *KCO* mice. ITR = internal tandem repeat for AAV2. CAGGS = CMV enhancer chicken beta-actin promoter. Leptin is murine cDNA. GFP is control. B) Schematic of AAV treatment for tumor interception experiment. AAV was administered at 6 weeks of age prior to development of significant tumor burden. Mice were analyzed 6 weeks later. Average percent change in body weight +/− s.e.m (n=8-10 mice/group) following AAV administration is shown. C) Representative histologic sections of pancreata of mice at endpoint demonstrate a reduction in CK19-positive ductal tumors with AAV-Leptin. Quantification of tumor burden (average % cross-sectional area of disease/total +/− s.e.m.) is shown (n=8-10 mice/group). D) Percent change in body weight (2 weeks post infection (p.i.)) from (C) plotted against tumor burden in (E) of AAV-Leptin-infected *KCO* mice. Each point represents one mouse (n=10). Greater weight loss is associated with less tumor burden. R^2^ and p-values are shown for linear regression E) Schematic for caloric restriction (CR) experiment. Mice began caloric restriction (1g/mouse/day of food) at 6 weeks of age as in (B). Controls remained on ad libitum diet (AL). Average percent change in body weight +/− s.e.m (n=6 mice/group) following intervention is shown. F) Representative histologic sections of pancreata of mice at endpoint demonstrate a reduction in CK19-positive ductal tumors with caloric restriction. Quantification of tumor burden (average % cross-sectional area of disease/total +/− s.e.m.) is shown (n=6 mice/group). p<0.05, **p<0.01, ***p<0.001, ****p<0.0001, two-tailed student’s t-test for all pairwise comparisons. Scale bars: 200 µm.

### Obesity promotes PDAC progression independent of new mutations

We next sought to identify the mechanisms by which obesity contributes to pancreatic tumorigenesis in this model. We postulated that obesity might induce DNA damage in the setting of enhanced oxidative stress (Usman and Volpi, 2018) and that tumor progression in *KCO* mice would depend on additional driver gene mutations beyond oncogenic *Kras*. Precedent for this hypothesis is provided by acceleration of oncogenic *Kras*-induced pancreatic tumor development in mice following inactivating mutations in *Trp53*, *Cdkn2a*, and *Smad4* (Bardeesy et al., 2006a; Bardeesy et al., 2006b; Hingorani et al., 2005; Muzumdar et al., 2016) – the hallmark recurrently mutated tumor suppressor genes (TSGs) in human PDAC (Bailey et al., 2016; The Cancer Genome Atlas Research Network, 2017)). In further support, survival of *KCO* mice was comparable to what we previously observed (Muzumdar et al., 2016) in *KPC* mice (*Pdx1-Cre; Kras^LSL-G12D/WT^; p53^flox/WT^* or *Pdx1-Cre; Kras^LSL-G12D/WT^; p53^LSL-R172H/WT^*) harboring heterozygous mutations in *p53* (p=0.93, log-rank test). Surprisingly, however, immunohistochemistry (IHC) for the hallmark TSGs revealed persistent protein expression, and exome sequencing confirmed the absence of inactivating point mutations in these genes (**Figure 3** and **Table S1**). Furthermore, we did not observe recurrent pathogenic mutations in other known PDAC (Bailey et al., 2016; The Cancer Genome Atlas Research Network, 2017) or pan-cancer driver genes (Bailey et al., 2018) in *KCO* tumors (**Tables S2-S3**). Contrary to our expectations, these data argue that additional driver gene mutations are not required for obesity to promote pancreatic cancer development. To determine if a similar phenomenon is evident in human tumors, we performed targeted exome sequencing and IHC to evaluate alterations of *KRAS*, *TP53*, *CDKN2A/p16*, and *SMAD4* in 184 human tumors with known patient BMI at diagnosis (**Table S4**). Although we observed a significant increase in *SMAD4* loss with elevated BMI (odds ratio 3.82 for BMI>30 kg/m^2^ compared to BMI<25 kg/m^2^), the total number of driver gene alterations per tumor, which we have previously shown predicts worse PDAC outcome (Qian et al., 2018), did not significantly differ across categories of BMI (**Table S5**). Together, these data suggest that obesity may function independently of driver gene alterations in both mice and humans to promote pancreatic tumorigenesis.

**Figure 3.**
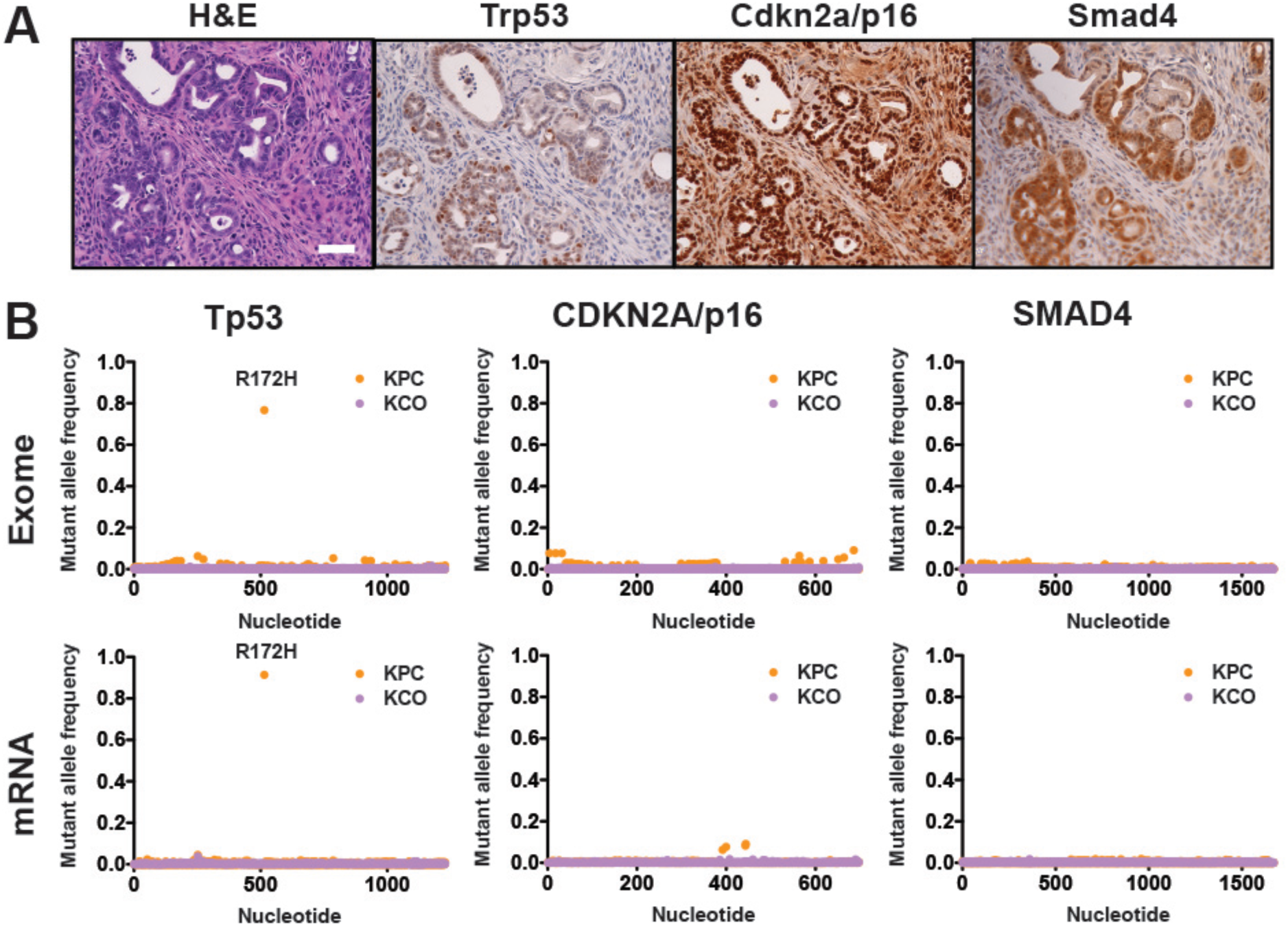
Absence of additional PDAC driver alterations in *KCO* tumors. A) Tumor suppressor genes frequently mutated in human cancers (*Trp53*, *Cdkn2A*/*p16*, and *Smad44*) were retained in *KCO* tumors as assessed by immunohistochemistry (IHC) and confirmed in 8/8 (100%) *KCO* tumors. Scale bar: 50 µm. B) Exome and mRNA sequencing did not reveal new single nucleotide variants (SNVs) in *Trp53* (murine *p53* gene), *Cdkn2A/p16*, or *Smad4* in *KCO* tumors. Average mutant allele fraction identified for each base in the coding sequence of each tumor suppressor gene is shown for *KCO* mice. A single *KPC* tumor sequenced in parallel shows the presence of the expected R172H codon change and serves as a positive control for these analyses.

### Enhanced tumor-associated inflammation and fibrosis in obesity

To elucidate potential non-mutational mechanisms of PDAC progression, we performed bulk RNA-sequencing (RNA-seq) on pancreata from obese *KCO* and non-obese *KC* and *KPC* mice exhibiting comparable tumor burden (**Table S6**), permitting analyses of both tumor cells and their microenvironment in cancer pathogenesis. Using Independent Component Analysis (ICA), an unsupervised blind source separation technique (Hyvarinen and Oja, 2000; Muzumdar et al., 2017), we identified gene expression signatures associated with inflammation and fibrosis as upregulated in *KCO* compared to *KC*/*KPC* mice (**Figures 4A-D** and **Tables S7-S8**), likely representing obesity-associated alterations in the tumor microenvironment. We confirmed abundant extracellular matrix deposition and immune cell infiltration by histologic analyses of *KCO* tumor sections (**Figure 4E**). To determine human relevance, we ranked primary human PDAC tumors from The Cancer Genome Atlas (TCGA) (The Cancer Genome Atlas Research Network, 2017) based on gene expression correlation to these *KCO* signatures. We classified tumors using previously-defined molecular subtypes (Bailey et al., 2016) and found the *KCO*-correlated tumors to be significantly enriched with the immunogenic subtype (**Figure 4F**), which is similarly associated with immune cell infiltration (Bailey et al., 2016; The Cancer Genome Atlas Research Network, 2017). In accordance with prior work using HFD models (Incio et al., 2016; Rooman et al., 2017), these data revealed that alterations in the fibroinflammatory microenvironment are a major feature of obesity-associated pancreatic tumors. Nonetheless, we observed no effect of anti-inflammatory (aspirin) or anti-fibrotic (metformin) (Chen et al., 2017; Rangarajan et al., 2018) drugs – shown to mitigate pancreatic cancer risk in selected (Li et al., 2009; Zhang et al., 2015) but not all patient cohorts (Khalaf et al., 2018) – on tumor progression in *KCO* mice (**Figure S3**).

**Figure 4.**
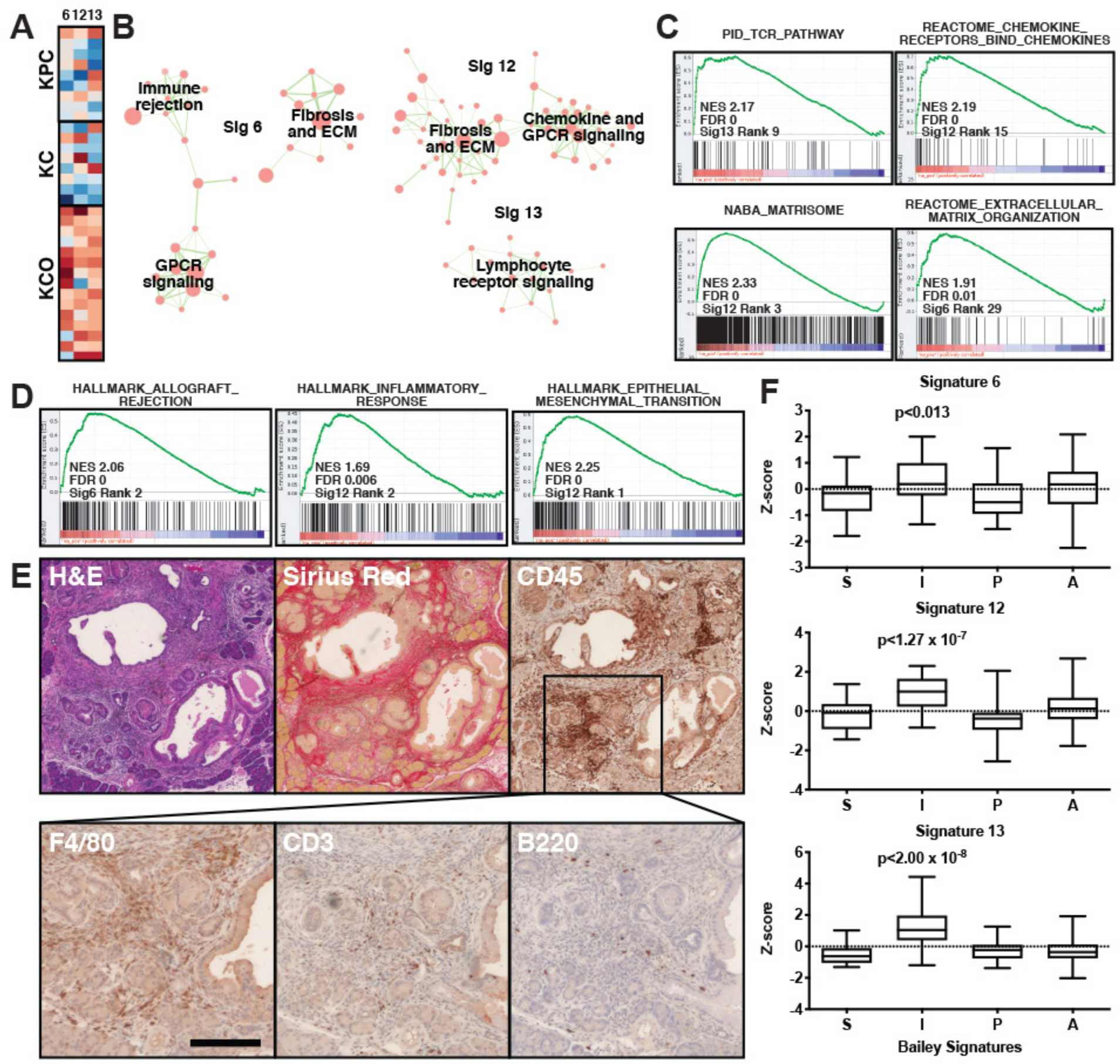
Enhanced inflammation and fibrosis in *KCO* tumors. A) Heat map of row normalized Z-scores (of mixing weights from the ICA decomposition) for ICA signatures (columns) showing statistically significant differences (p<0.01, Mann-Whitney test) between *KCO* mice versus non-obese *KC* and *KPC* mice. Signature numbers (6, 12, 13) are denoted above. Rows are individual tumors where the Z-score reflects relative correspondence between a tumor and the expression signature pattern (degree of positive or negative correlation). Red corresponds to positive and blue to negative Z-scores. B) Network representation of overlapping enriched GSEA/MSigDB gene sets associated with the signatures in (a). Cellular processes associated with related gene sets are listed. C) GSEA using the curated gene set collection (C2 in MSigDB) revealed an association between *KCO* tumors and genes involved in immune cell activation/signaling and extracellular matrix (ECM)/fibrosis. Normalized enrichment score (NES), false discovery rate (FDR), ICA signature (*KCO* vs. *KC*/*KPC*), and MSigDB gene set rank (for C2 sets) used in analysis are shown. See **Table S8** for complete list. D) GSEA showed an association between *KCO* tumors and inflammation and fibrosis genes in comparison to hallmark genes set collection (H in MSigDB). E) Histologic analyses revealed extensive fibrosis (Sirius red staining of collagen) and CD45+ immune cell infiltration predominantly with F4/80+ macrophages and sparse presence of CD3+ T and B220+ B cells. Scale bars: 100 µm F) Box and whisker plots (boxes denote 25^th^-75^th^ percentile, error bars denote min/max) of standardized signature scores of TCGA (scored using ssGSEA (Barbie et al., 2009) for expression correlation with each of the signatures in (a) (6, 12, and 13) for each molecular subtype (S = squamous (n=31), I = immunogenic (n=28), P = progenitor (n=53), A = ADEX (n=38)). p-values confirmed significant enrichment of highly-correlated tumors with the immunogenic subtype (hypergeometric test) for each signature.

### Pancreatic islet cell adaptations in tumors of obese mice

One of the most notable results from our bulk RNA-seq was significant upregulation of genes associated with pancreatic islet cell function in *KCO* compared to *KC/KPC* pancreata (**Figures 5A**). Although expression levels of general neuroendocrine markers typically used to identify islets in pancreatic tissue were unchanged or only mildly elevated in *KCO* pancreata (**Figure 5B**), we observed a marked increase in the expression of genes encoding islet hormones, proteolytic enzymes involved in islet prohormone cleavage, and secretory granule proteins (**Figure 5C**). We confirmed the gene expression data by IHC on islets in tumor-bearing mice. While glucagon (GCG) and its proteolytic product GLP-1 are expressed in peripherally-located alpha islet cells in non-obese rodents (Barreto et al., 2010), *KCO* mice displayed an expansion of GCG and GLP-1 production throughout islets (**Figure 5D**). These observations are consistent with prior research demonstrating increased GLP-1 secretion from islets of obese humans and mice (Linnemann et al., 2015). Together, our findings support islet adaptation to enhance hormone production, processing, and secretion in the setting of obesity.

**Figure 5.**
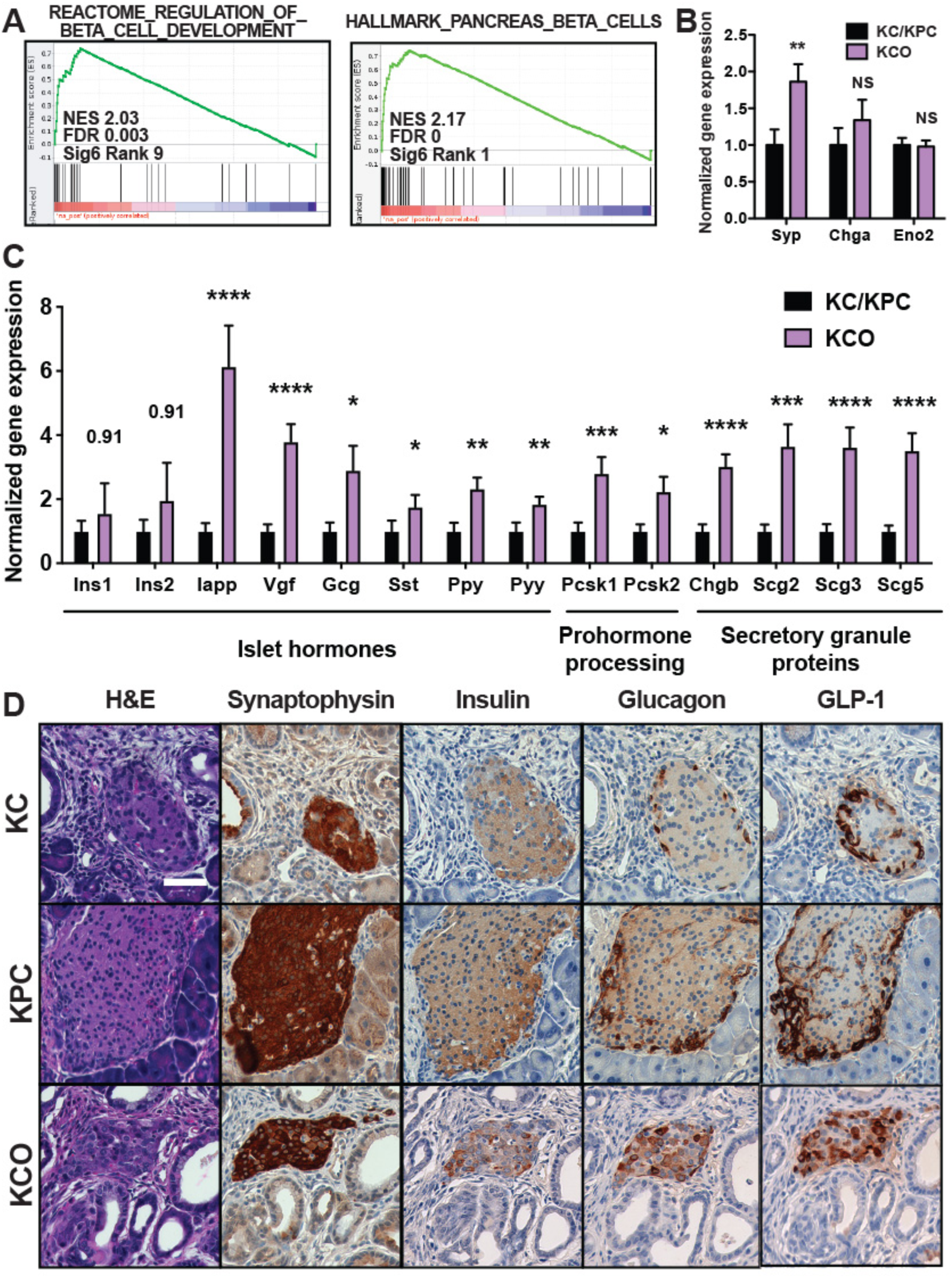
Pancreatic islet adaptation in *KCO* tumors. A) Gene set enrichment analysis (GSEA) using the curated (C2 in MSigDB) and hallmark (H in MSigDB).gene set collections revealed an association between *KCO* tumors and genes expressed in pancreatic beta cells. Normalized enrichment score (NES), false discovery rate (FDR), MSigDB gene set rank (for C2 and H sets), and the ICA signature (*KCO* vs. *KC*/*KPC*) used in the analyses are shown. B) Normalized gene expression (mean RNA-seq expression counts +/− s.e.m. normalized to non-obese *KC/KPC* tumors) for general neuroendocrine markers observed in pancreatic islet cells (synaptophysin (*Syp*), chromogranin A (*Chga*), and neuron-specific enolase (*Eno2*)), showed mild to no significant difference between *KCO* (n=15) and *KC/KPC* (n=17) mice. C) Normalized expression (mean RNA-seq expression counts +/− s.e.m. normalized to non-obese models) of islet hormones (glucagon (*Gcg*), islet amyloid polypeptide (*Iapp*), *Vgf*, somatostatin (*Sst*), *Ppy*, and *Pyy*), prohormone endopeptidases (*Pcsk1*, *Pcsk2*), and secretory granule proteins *Chgb*, *Scg2*, *Scg3*, and *Scg5*) are upregulated in *KCO* mice compared to non-obese models. * p<0.05, **p<0.01, ***p<0.001, ****p<0.0001, two-tailed Mann-Whitney test, comparing *KCO to KC/KPC* expression for each gene. D) Immunohistochemistry showed aberrant glucagon and GLP-1 expression throughout pancreatic islets in *KCO* mice compared to non-obese controls, consistent with upregulation of *Gcg* and *Pcsk1*/*Pcsk2* observed by RNA-seq in (C). Scale bar: 50 µm.

Given these islet adaptations, we postulated that enhanced insulin secretion may promote local pancreatic tumorigenesis in obese mice. Although aberrant insulin signaling is commonly proposed as a mechanism by which obesity drives tumorigenesis, direct *in vivo* evidence, especially in PDAC, is limited (Wang et al., 2018; Zhang et al., 2019). As expected, *ob/ob* mice exhibited systemic hyperinsulinemia with increased fasting serum insulin (**Figure S4A**). In contrast, glucose-stimulated insulin secretion by primary islets derived from *ob/ob* mice was reduced (**Figures S4B-D**), arguing that local insulin secretion is severely impaired. Furthermore, analysis of insulin receptor phosphorylation by IHC suggested comparable signaling activation in tumor cells from both non-obese *KC* and obese *KCO* mice (**Figure S4E**). These data argue against local increases in insulin or insulin signaling as a primary driver of pancreatic tumorigenesis, suggesting that additional islet-derived secreted factors may be involved.

### Single cell RNA-sequencing identifies beta cell expression of CCK in obesity

To identify such hormones at greater resolution, we performed single cell RNA-sequencing (scRNA-seq) on islets isolated from congenic C57/B6 wild-type and *ob/ob* mice. Visualization using PHATE (Moon et al., 2019), a method for dimensionality reduction of single-cell data, displayed clusters corresponding to single hormone-expressing beta (insulin), alpha (glucagon), delta (somatostatin), and PP (Ppy) cells (**Figure 6A**). Within these cell types, there was only moderate overlap in the beta cell populations comparing *ob/ob* and wild-type mice (**Figure 6B**), consistent with significant adaptive changes in gene expression to obesity. Moreover, *ob/ob* islets exhibited a higher proportion of beta cells and a commensurate reduction in alpha, delta, and PP cells (**Figure 6B**). Beta cells from *ob/ob* mice showed increased expression of genes involved in protein translation, endoplasmic reticulum (ER) stress, and the secretory pathway, consistent with enhanced synthesis of secreted proteins (**Figure 6C** and **Table S9**). These genes included hormones and secretory granule proteins also observed in *KCO* mice (**Figure 6D**), confirming results from bulk tumor RNA-sequencing.

**Figure 6.**
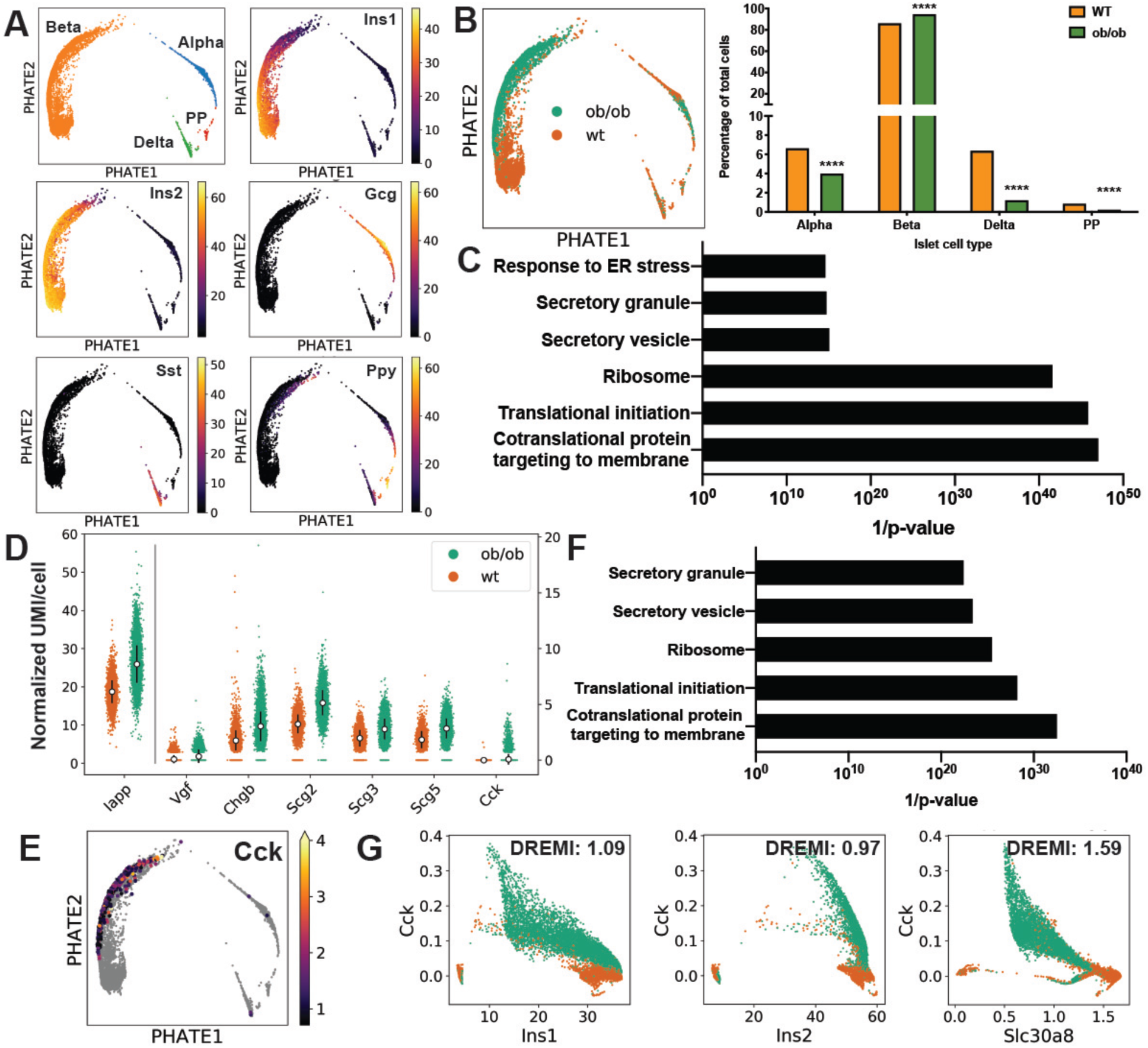
scRNA-seq identifies beta cell expression of CCK in obesity. A) PHATE visualization plots for single-hormone expressing islet clusters identifies *Ins1*/*Ins2*-expressing beta cells, *Gcg*-expressing alpha cells, *Sst*-expressing delta cells, and *Ppy*-expressing PP cells amongst all sequenced cells. B) PHATE visualization plot overlaying cells sequenced from wild-type (WT) and *ob/ob* mice on clusters in (A) shows only moderate overlap in beta and alpha clusters. The number of cells in each cluster were also statistically different between genotypes (****p<0.0001, χ-square). C) Gene set enrichment analysis (GSEA) using gene ontology (C5 GO) gene sets from MSigDB for genes upregulated in *ob/ob* versus WT beta cells (q<0.0001, log2FC>0.5, mean normalized expression counts >0.5). D) Mean single cell expression counts (square-root transformed library size-normalized UMI/cell +/− s.d.) for beta cells shows upregulation of hormones (*Iapp*, *Vgf*, and *Cck*) and secretory granule proteins (*Chgb*, *Scg2*, *Scg3*, and *Scg5*) in *ob/ob* islets. E) PHATE visualization shows Cck expression exclusively in beta cells. Cck is expressed in *ob/ob*, but not WT, islets as shown in (D). F) GSEA using gene ontology (C5 GO) gene sets from MSigDB for genes upregulated in CCK-positive (expression count > 0) versus CCK-negative *ob/ob* beta cells (q<0.0001, mean normalized expression counts > Cck). G) Gene-gene expression plots after MAGIC (van Dijk et al., 2018) shows an inverse relationship between the expression of *Ins1*, *Ins2*, and *Slc30a8* with *Cck*. *k*NN-DREMI scores showing the strength of the association between gene-pairs are listed.

The top differentially expressed gene (by fold change) comparing *ob/ob* and wild-type beta cells was the gut hormone cholecystokinin (CCK) (**Figure 6D** and **Table S9**). CCK is normally synthesized and secreted by enteroendocrine cells of the duodenum to induce pancreatic acinar cell zymogen release for digestion. Its dysregulation in islets of obese mice (HFD and *ob/ob*) has been reported to promote islet cell survival in the setting of insulin resistance, toxins, or other stresses (Lavine et al., 2015; Lavine et al., 2010). We confirmed islet CCK expression by IHC in *ob/ob* mice (**Figures S5A**). We further observed CCK expression exclusively in a subset of beta cells (**Figure 6E**), those of which exhibited enhanced expression of protein synthesis and secretory granule genes than their CCK-negative counterparts (**Figure 6F** and **Table S8**). Furthermore, CCK expression was inversely associated with the expression of insulin genes (*Ins1* and *Ins2*) and *Slc30a8*, a zinc transporter involved in insulin granule maturation (**Figure 6G**), arguing that CCK – rather than insulin – secretion may predominate in these cells.

### Local islet CCK signaling drives pancreatic ductal tumorigenesis

Since exogenous administration of cerulein, a CCK analogue, promotes *Kras*-driven pancreatic tumorigenesis in mice (Guerra et al., 2007), we hypothesized that, in the setting of obesity, endogenous islet-derived CCK could drive pancreatic cancer development in the local microenvironment. Indeed, systemic CCK expression was not increased in obese mice (**Figure S5B**), arguing that islet CCK would principally act locally. In *KCO* mice, CCK expression was specifically observed in islets and markedly reduced by weight loss interventions (**Figures 7A-B** and **S5C-D**), consistent with an adaptive response to obesity. To determine human relevance, we performed IHC on pancreata from PDAC patients and observed islet expression of CCK in ∼60% of samples (**Figures 7C-D**). Although CCK expression did not correlate with BMI (**Figure 7D**), cancer-associated weight loss prior to diagnosis confounds this analysis. Therefore, we analyzed islets obtained from human donors without known malignancy and observed a positive relationship between BMI and CCK expression (**Figure 7E**). Together, these data demonstrate that obesity promotes islet CCK expression.

**Figure 7.**
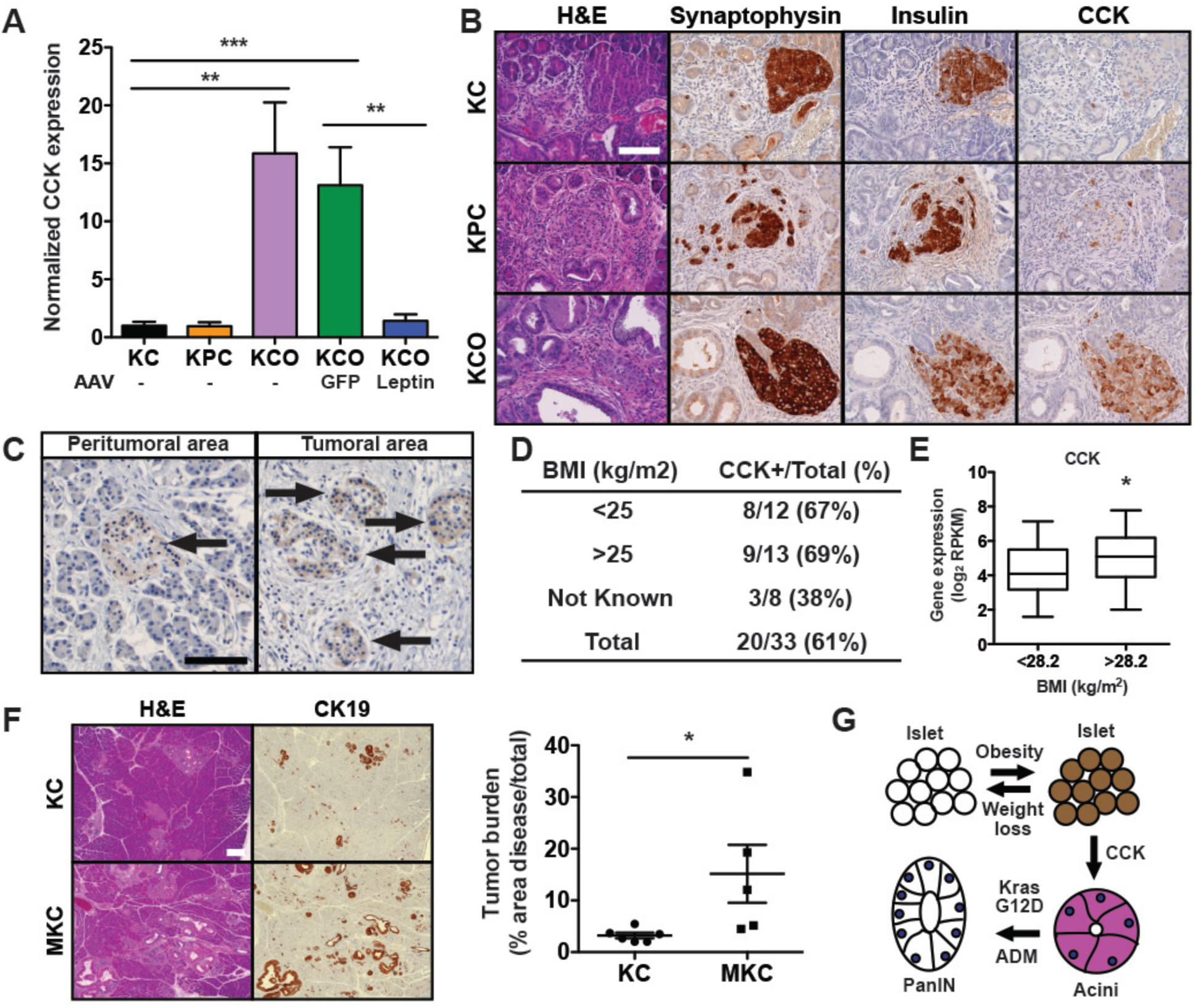
Islet-derived cholecystokinin promotes pancreatic ductal cancer development. A) Normalized expression (mean RNA-seq expression counts +/− s.e.m. normalized to *KC* tumors) of cholecystokinin (CCK) in obese (*KCO* and *KCO* mice treated with AAV-GFP) and non-obese models (*KC*, *KPC*, and *KCO* mice treated with AAV-Leptin) (n=5-9 per group). **p<0.01, ***p<0.001, two-tailed Mann-Whitney test. B) CCK was aberrantly overexpressed in pancreatic islets (synaptophysin-positive) of *KCO* compared to *KC* and *KPC* mice, including in insulin-producing cells. Scale bar: 100 µm. C) CCK was expressed specifically in islet cells (arrows) in human PDAC by immunohistochemistry (IHC) in intra-tumoral and peri-tumoral areas. Scale bar: 100 µm. D) The majority of human PDAC specimens displayed islet expression by IHC at all BMI levels (<25 kg/m^2^ (normal) and >25 kg/m^2^ (overweight or obese)). E) CCK expression in isolated pancreatic islets from human donors with high BMI (n=53, above median US BMI of 28.2 kg/m^2^) was significantly (*p<0.05, two-tailed Mann-Whitney test) greater than those from low BMI donors (n=55, below median). Box and whisker plots (boxes denote 25^th^-75^th^ percentile, error bars denote min/max) are shown. F) Representative histologic sections of pancreata demonstrate an increase in tumorigenesis in *MKC* (*MIP-CCK; KC*) mice compared to *KC* mice. Quantification of tumor burden (average % cross-sectional area of disease/total +/− s.e.m.) in *KC* and *MKC* mice at 3 months (n=5-6 mice/group). * p<0.05, two-tailed student’s t-test. Scale bar: 200 µm. G) Model for obesity-associated islet adaptations in PDAC development. Brown denotes CCK expression. ADM = acinar-to-ductal metaplasia.

To test directly whether islet CCK contributes to tumor progression, we utilized *MIP-CCK* transgenic mice (Lavine et al., 2015), which exhibit pancreatic islet-specific CCK expression (**Figure S6A**) comparable to *ob/ob* mice without systemic CCK elevation (Lavine et al., 2015). Remarkably, non-obese compound transgenic mice (*MKC*: *MIP-CCK; Pdx1-Cre; Kras^LSL-G12D/WT^*) displayed a significant increase in tumor burden compared to *KC* controls, (**Figures 7F**), supporting the hypothesis that islet CCK overexpression functions as an independent driver of pancreatic ductal tumorigenesis.

Previous work showed that the non-selective competitive CCKR antagonist proglumide at doses that block physiologic CCK function impedes tumor progression in non-obese *KC* mice (Smith et al., 2014). Since obesity-augmented islet CCK expression appears to function as a local driver of early PDAC progression, we anticipated that proglumide treatment should be insufficient to impair progression in the setting of supraphysiologic local concentrations of CCK in *KCO* mice. Indeed, although proglumide induced the expected systemic pharmacodynamic effect of slowing weight gain, no significant impact on tumor development was observed in our model (**Figure S3**). Thus, these data provide evidence for the importance of aberrantly-expressed pancreatic endocrine hormones beyond insulin in driving obesity-associated PDAC.

### Endocrine-exocrine CCK signaling in obesity

Finally, we explored the mechanism by which islet CCK functions in the local microenvironment to drive tumor development. PDAC is commonly thought to arise from acinar or ductal cells (Kopp et al., 2012; Ray et al., 2011), suggesting that the exocrine pancreas is the target for islet CCK. Evidence supports a direct effect of CCK on CCK receptor signaling in rodent acinar cells. Although CCK receptor expression in human acinar cells is lower than observed in rodents, recent work supports a similar direct effect by physiologic and supraphysiologic concentrations of CCK on intracellular signaling and zymogen release in primary human acinar cells (Liang et al., 2017; Murphy et al., 2008). Alternatively, CCK has been shown to stimulate insulin secretion from beta cells (Lo et al., 2011), which in turn could modulate early tumorigenesis. Contrary to this latter hypothesis, CCK receptor (*Cckar* or *Cckbr*) transcripts were detectable in <0.05% of beta cells (compared to 12% of Prss2+/Cela1+ acinar cells) profiled from wild-type and *ob/ob* mice by scRNA-seq at our depth of sequencing. Furthermore, *MIP-CCK* mice do not exhibit increased insulin release during fasting or following glucose stimulation (Lavine et al., 2015), and *ob/ob* islets were impaired in glucose-stimulated insulin secretion (**Figure S4B-D**). Therefore, we conclude that endocrine-derived CCK acts on acinar cells to drive tumorigenesis.

While exogenous cerulein administration is commonly used as a model for experimental pancreatitis (Guerra et al., 2007), our data support a pro-tumorigenic effect of islet CCK independent of inflammation. *MIP-CCK* mice did not exhibit overt evidence of inflammation or fibrosis (**Figure S6A**) even up to one year of age (Lavine et al., 2015). Conversely, acute (cerulein- or arginine-induced) or chronic (cerulein-induced) pancreatitis did not elicit islet CCK expression in mice (**Figures S6B-C**). CCK expression was similarly not significantly observed in islets of patients with chronic pancreatitis (**Figure S6D**). Since, in explant cultures, cerulein can stimulate acinar-to-ductal metaplasia (ADM) (Ardito et al., 2012), a prerequisite step in acinar-derived PDAC development (Kopp et al., 2012), we propose that islet CCK cooperates with oncogenic *Kras* to drive ADM and early tumorigenesis (**Figure 7G**). Importantly, islet CCK expression is reversible, supporting the role of anti-obesity strategies in PDAC prevention.

## DISCUSSION

In this study, we leverage an autochthonous mouse PDAC model and demonstrate that obesity plays a causal and reversible role in early pancreatic tumor formation and progression to advanced disease. We define microenvironmental rather than intrinsic genetic alterations as hallmarks of tumorigenesis in obesity, including increased inflammation, fibrosis, and pancreatic islet cell adaptation, the latter of which we show can drive tumorigenesis. Specifically, we observe aberrant islet expression of the neuropeptide CCK and a pro-tumorigenic role for islet-derived CCK in PDAC development. Together, these data support a unifying mechanism linking obesity, changes in the local microenvironment, and tumorigenesis via adaptive endocrine-exocrine signaling by pro-tumorigenic hormones.

Hormones have long been postulated as contributing to obesity-associated cancers with principal mechanisms including dysregulation of 1) islet-derived insulin or insulin-like growth factor 1 (IGF1); 2) adipocyte secreted factors (adipokines) including leptin and adiponectin; and 3) steroid sex hormones (Murphy et al., 2018; Ulrich et al., 2018). Of these mechanisms, epidemiologic studies support a role for both insulin/IGF1 and adipokines in pancreatic cancer development, as elevated levels of insulin, proinsulin, and leptin and low levels of IGF binding protein-1 and adiponectin are associated with increased PDAC risk (Babic et al., 2016; Bao et al., 2013a; Bao et al., 2013b; Wolpin et al., 2013; Wolpin et al., 2007). Though previous studies in cell lines and xenografts have demonstrated that leptin can promote PDAC proliferation (Harbuzariu et al., 2017; Mendonsa et al., 2015), our observation that obese mice deficient in leptin signaling (*ob/ob* and *db/db*) exhibit enhanced *Kras*-driven pancreatic tumor progression supports a leptin-independent mechanism for obesity in tumor promotion. Furthermore, the capacity for caloric restriction-induced weight loss to intercept progression in *KCO* mice is consistent with obesity, rather than dysregulation of leptin signaling or the associated adaptive physiologic response to starvation (Friedman, 2019), as a driver of PDAC development.

Apart from insulin (Wang et al., 2018; Zhang et al., 2019), evidence for a pro-tumorigenic role of pancreatic islet hormones in cancer development has been lacking until now. Our study shows that obesity-associated aberrant islet CCK expression drives cancer development in mice through a local effect in the pancreas. Ultrastructural and perfusion studies endorse the existence of an islet portal system perfusing exocrine cells (the islet-acinar axis) (Barreto et al., 2010; Williams and Goldfine, 1985). Alternatively, capillary leak or neovascularization in the setting of islet cell expansion and local tissue injury from early developing tumors may alter blood flow and permit high concentrations of islet-derived CCK to access exocrine cells. The significance of obesity-driven islet adaptations is highlighted by the observation that CCK-expressing beta cells show diminished expression of genes involved in insulin secretion, including insulin itself, consistent with major alterations in the secretome. We further demonstrate that CCK is expressed in islets in human PDAC biospecimens and that islet CCK expression correlates with BMI, supporting the possibility that a similar phenomenon occurs in human PDAC development. Our results add to the growing body of evidence that obesity may be pro-tumorigenic through its microenvironmental rather than systemic consequences (Olson et al., 2017). Although evidence for a role for CCK signaling in PDAC progression exists (Nadella et al., 2018; Smith et al., 2014; Smith and Solomon, 2014), prior studies focused on intestinal-derived CCK, the physiologic source of systemic CCK. On the contrary, our work highlights a local source of CCK – induced by obesity within the pancreas itself – and offers the first example of endocrine-exocrine signaling beyond insulin in PDAC development.

Finally, our results demonstrate that pro-tumorigenic hormonal adaptations, including islet CCK expression, are reversible with weight loss, arguing for the use of anti-obesity strategies for PDAC prevention. Medical and surgical interventions to promote weight loss in human populations have not been adequately powered to study their capacity to alter PDAC development. Nonetheless, weight loss due to bariatric surgery reduces overall cancer incidence (Xu et al., 2018). Our data endorse an effect of weight loss on early PDAC progression, revealing a temporal window in which anti-obesity strategies may prevent or intercept cancer.

## ACKNOWLEDGEMENTS

We thank F. Gorelick, M. Lemmon, and F. Wilson for critical reading of the manuscript; R. Perry for helpful discussions; K. Mercer for technical assistance; K. Cormier and C. Condon from the Hope Babette Tang (1983) Histology Facility and A. Brooks, P. Gaule, Y. Bai, B. Acs, and D. Rhim from Yale Pathology Tissue Services for histology assistance; M. Robert and R. Bronson for pathology assistance; S. Levine, V. Dayal, and J. Penterman from the MIT BioMicro Center for bulk RNA-seq support; J. Heitke, G. Wang, C. Castaldi, S. Mane from the Yale Center for Genome Analysis (YCGA) for assisting with scRNA-seq; the University of Iowa Viral Vector core for AAV constructs and generation; M. Batsu and R. Jacobs from the Yale Diabetes Research Core for serum ELISA/RIA studies; the Oxford Human Islet Isolation facility for the provision of human islets for research; T. Kolodecik and F. Gorelick for providing murine pancreatitis specimens; F. Zheng for CRISPR constructs; and A. Berns, D. B. Davis, A. Lowy, L. Luo, and Jackson Laboratories for mice.

This work was supported in part by a Lustgarten Foundation Research Investigator Grant (C.S.F. and M.D.M.), Cancer Center Support (core) grants P30-CA016359 (C.S.F.) and P30-CA014051 (T.J.) from the National Cancer Institute, a Pilot Grant from the Yale Cancer Center (M.D.M.), and a Yale Cancer Innovations Award (M.D.M.). The National Institute for Health Research, Oxford Biomedical Research Centre funded islet provision at the Oxford Human Islet Isolation facility. C.G. is funded by an NIH/NIGMS training grants (T32-GM007499). D.B.B. is funded by the NICHD (F31-HD097958). A.B. is funded by an NCI Mentored Cancer Prevention, Control, Behavioral Sciences, and Population Sciences Career Development Award (K07CA222159) and a Bob Parsons Fellowship. A.L.G. is a Wellcome Senior Fellow in Basic Biomedical Science. Her work was funded in Oxford by the Wellcome Trust (095101, 200837, 106130, 203141, Medical Research Council (MR/L020149/1), European Union Horizon 2020 Programme (T2D Systems), and NIH (U01-DK105535; U01-DK085545). M.I.M. is a Wellcome Senior Investigator and an NIHR Senior Investigator. Relevant funding support for this work comes from Wellcome (090532, 106130, 098381, 203141, 212259), NIDDK (U01-DK105535), and NIHR (NF-SI-0617-10090). The views expressed in this article are those of the author(s) and not necessarily those of the NHS, the NIHR, or the Department of Health. R.G.K. is funded by grants P30DK045735 and R01DK110181 from the NIDDK and a Yale Cancer Innovations Award. S.K. is funded by the NIH (R01-GM130847), HIPC (2U19AI089992), and the Chan Zuckerberg Initiative (CZF2019-002440). B.M.W. acknowledges support from the Hale Family Center for Pancreatic Cancer Research, Lustgarten Foundation, NIH U01-CA210171, Stand Up to Cancer, Noble Effort Fund, Wexler Family Fund, and Promises for Purple. T.J. is a Howard Hughes Medical Institute Investigator, the David H. Koch Professor of Biology, and a Daniel K. Ludwig Scholar. M.D.M. is supported by an NCI Mentored Clinical Scientist Research Career Development Award (K08CA2080016) and was supported by a KL2/Catalyst Medical Research Investigator Training award (an appointed KL2 award) from Harvard Catalyst | The Harvard Clinical and Translational Science Center (National Center for Research Resources and the National Center for Advancing Translational Sciences) (KL2 TR001100), a Conquer Cancer Foundation-American Society for Clinical Oncology (CCF-ASCO) Young Investigator Award, a Career Development Award from the Dana-Farber Cancer Institute/Harvard Cancer Center SPORE in Gastrointestinal Cancer, and the NIH Loan Repayment Program. The content is solely the responsibility of the authors and does not necessarily represent the official views of the National Institutes of Health.

## AUTHOR CONTRIBUTIONS

T.J., C.S.F., and M.D.M. designed and supervised the study; K.M.C., J.S., L.L., K.J.D., C.G., R.R., and M.D.M. performed experiments; D.B.B., A.B., and S.K. conducted computational analyses; R.C., X.Z., and R.G.K. performed and analyzed islet isolation and perifusion studies; A.B., J.N., S.A.V., A.D.C., and B.M.W. performed targeted exome and IHC analyses on human PDAC; D.T.C., R.F.D., A.F.H., A.C.K., J.J.W., and M.D.B. provided human PDAC and pancreatitis biospecimens; V.N., A.L.G., and M.I.M. acquired and analyzed human donor islet gene expression data; M.D.M. wrote the manuscript with comments from all authors.

## METHODS

### Animals studies

Animal studies were approved by the Massachusetts Institute of Technology (MIT) and Yale University Institutional Animal Care and Use Committees (IACUC). *ob* (Stock #000632), *db* (Stock #000697), *Pdx1-Cre* (Stock #014647), *p53^fl/fl^* (Stock #008462), *MADM11-GT* (Stock #013749), and *MADM11-TG* (Stock #013751) mice were obtained from the Jackson Laboratory. *MIP-CCK* mice (Lavine et al., 2015) were obtained from D. B. Davis of the University of Wisconsin. *LSL-Kras^G12D^*, *p53^R172H/WT^*, *p53^KO/WT^*, and *Kras^LA2^* mice were generated previously by the Jacks lab (Jacks et al., 1994; Jackson et al., 2001; Johnson et al., 2001; Olive et al., 2004). Animals of both genders were used in all experiments. Mice were housed in a specific-pathogen free facility, kept at room temperature with standard day-night cycles, and maintained on a mixed background unless otherwise specified.

*Pdx1-Cre; Kras^LSL-G12D/WT^(KC), KC; ob/+* and *KC; ob/ob* (*KCO*), *KC; db/+*, *KC; db/db*, *Kras^LA2^; +/+*, and *Kras^LA2^; ob/ob* were produced by intercrossing mice with the appropriate alleles. *KPC* mice, including *KC; p53^R172H/WT^*; *KC; p53^fl/^*^WT^, *KC; p53^R172H/fl^, KC; p53^fl/fl^*, and *KC; MADM11-GT,p53^WT^/MADM11-TG-p53^KO^* (*MADM-p53)*, were generated as described previously (Muzumdar et al., 2016). Mice were genotyped using tail DNA by the HotShot method. Genotyping primers and protocols have been previously reported (Ellett et al., 2009; Lavine et al., 2015; Muzumdar et al., 2016) or are available online (http://www.jax.org).

Animals were monitored at least weekly for signs of morbidity and were euthanized by CO2 asphyxiation when they met euthanasia criteria. Date of euthanasia relative to birth date was used for Kaplan-Meier survival analyses. Weights were monitored using a standard small animal scale. Glucose was measured on a OneTouch Ultra2 glucometer on whole blood collected by tail clip. For serum studies, whole blood was collected from anesthetized mice by retro-orbital bleed or terminal cardiac puncture, allowed to clot for 30 minutes at room temperature, and spun through a serum separator tube (Sarstedt) at 10,000 rpm for 5 min prior to storage at −80°C or analysis. Serum CCK was measured by EIA (RayBiotech, Cat# EIAM-CCK). Serum insulin (RIA, Linco Cat# RI-I3K), C-peptide (ELISA, ALPCO Cat# 80-CPTRT-E01), corticosterone (RIA, MP Biomedicals, LLC), free fatty acids (FFA), and triglycerides (TG) were measured by the Yale Diabetes Research Core.

### Mouse treatment studies

For diet studies, *Kras^LA2^* mice were started on high-fat (60% kcal fat, Research Diets 12492) or low-fat (10% kcal fat, Research Diets 12240J) diet at 4 weeks of age. For drug studies, *KCO* mice were treated with pharmaceutical-grade aspirin (2 mg/mL, Spectrum Chemical Group), metformin (2 mg/mL, Spectrum Chemical Group), and proglumide (0.1 mg/mL, Sigma) in drinking water starting at 6 weeks of age and treated for 6 weeks continuously. Water consumption was measured weekly when treated water was replaced. *KCO* mice drank 7.69 +/− 0.58 (s.d.), 8.95 +/− 2.24, 8.09 +/− 3.21, 9.70 +/− 1.49 mL drinking water daily for untreated (control), aspirin, metformin, and proglumide, respectively. Water intake was not statistically different from control for each treatment condition (two-tailed student’s t-test). Based on water consumption, average daily doses were 17.9, 16.2, and 0.97 mg for aspirin, metformin, and proglumide, leading to an average dose exposure of ∼456, 436, and 26 mg/kg per day, respectively, in-line with prior studies using these agents (Chandel et al., 2016; Smith et al., 2014). Mice were randomized following pre-stratification for gender in all treatment conditions.

### Murine tissue preparation, histology, and quantification

Mice were euthanized by CO2 asphyxiation and tissue was dissected, fixed in 10% neutral-buffered formalin overnight, and dehydrated in 70% ethanol prior to paraffin-embedding. Adjacent 5 µm sections were cut and stained with hematoxylin and eosin or picosirius red or subject to immunohistochemistry (IHC). Primary antibodies for IHC are listed below. Mach 2 HRP-labeled anti-mouse, anti-rabbit, or anti-guinea pig micro-polymers (Biocare Medical) were used for primary antibody detection using a Thermo Scientific Autostainer 360. Slides were imaged with a modified Nikon T2R inverted microscope (MVI), 4x/10x/20x/40x objectives, and a 2.8 MP CoolSNAP Dyno CCD camera (Photometrics). Monochromatic red, green, and blue images were merged using ImageJ software (NIH). Percentage of tumor burden was quantified blinded to group on scanned slides (Aperio) by measuring cross-sectional area of tumor-bearing areas (PanIN or PDAC including stroma) relative to measured total pancreas area using ImageJ or QuPath v0.1.2 software.

### Adeno-associated virus (AAV) generation

Total RNA was isolated from flash frozen adipose tissue derived from *db/db* mice using Trizol (Ambion) and reverse transcribed with a High Capacity cDNA Reverse Transcription kit (Thermo Fisher Scientific). Murine leptin cDNA was PCR amplified with Primestar high-fidelity PCR mix (Takera) using primers (forward 5’-ATGTGCTGGAGACCCCTGT-3’, reverse 5’-TCAGCATTCAGGGCTAACATCCAACT-3’) synthesized by the Koch Institute Swanson Biotechnology Center. Amplified leptin cDNA was cloned into the XhoI and NotI sites of the AAV2 backbone vector (*pFBAAVCAGmcsBgHpa*) provided by the University of Iowa Viral Vector Core (UIVVC), which generated AAV2/1-Leptin using a baculovirus system. AAV2/1-GFP, cloned into the same AAV2 plasmid vector, was purchased from UIVVC. AAV-induced leptin expression was verified in 293T cells by western blotting 48 hours after infection. 1×10^11^ AAV viral particles were administered to mice via a single injection in the tibialis anterior muscle and circulating leptin levels were measured in the serum by ELISA (Crystal Chem Cat# 90030).

### Antibodies

| Antibody Target | Species | Company | Product Number | Application | Dilution |
| --- | --- | --- | --- | --- | --- |
| B220 | Rat | BD Biosciences | 550286 | IHC | 1:100 |
| Beta-tubulin | Rabbit | Cell Signaling Technology | 2128 | WB | 1:1000 |
| Cd3 | Rabbit | Abcam | ab16669 | IHC | 1:300 |
| Cd45 LCA | Rabbit | Abcam | ab10558 | IHC | 1:100-1:500 |
| Cdkn2a | Rabbit | Abcam | ab54210 | IHC | 1:500 |
| Cholecystokinin | Rabbit | Immunostar | 20078 | IHC | 1:4000-1:8000 |
| Ck19 | Rabbit | Abcam | ab133496 | IHC | 1:800 |
| Erk1/2 | Mouse | Cell Signaling Technology | 9107 | WB | 1:1000 |
| F4/80 | Rabbit | Invitrogen | ma5-16363 | IHC | 1:100 |
| Gfp | Rabbit | Cell Signaling Technology | 2956 | WB | 1:1000 |
| Hsp90 | Mouse | BD Biosciences | 610418 | WB | 1:10,000 |
| Insulin | Guinea Pig | Accurate Chemical | YNGPSWINS7S | IHC | 1:300 |
| Leptin (OB) | Rabbit | Santa Cruz Biotechnology | sc-842 | WB | 1:200 |
| p53 | Rabbit | Novacastra | NCL-p53-CM5p | IHC | 1:400 |
| pErk1/2 (T202/Y204) | Rabbit | Cell Signaling Technology | 4370 | WB | 1:1000 |
| pInsR/Igf1R (Y1162/Y1163) | Rabbit | Thermo Scientific | 44-804G | IHC | 1:100 |
| pStat3 (Y705) | Rabbit | Cell Signaling Technology | 9145 | WB | 1:1000 |
| Smad4 | Rabbit | Abcam | ab40759 | IHC | 1:2000 |
| Stat3 | Mouse | Cell Signaling Technology | 9139 | WB | 1:1000 |
| Synaptophysin | Rabbit | Neo Markers | RB-1461 | IHC | 1:100 |

### Murine PDAC cell line derivation, culture, and treatments

Murine *KPC* PDAC (mPDAC) cell lines (7307 and 7310) were isolated from primary autochthonous pancreatic tumors arising in congenic C57/B6 *KC; p53^R172H/WT^* mice (Hingorani et al., 2005). Briefly, tumors were enzymatically dissociated using a mix of collagenase IV, dispase, and DNAse I (Worthington) at 37°C for 30 minutes with further mechanical dissociation using a gentleMACS dissociator (Miltenyi Biotec). Cell suspensions were passed through a 100 µM filter, washed, and plated in complete media containing DMEM (Corning), 10% FBS (Thermo Fisher Scientific), and penicillin/streptomycin (Sigma). Cells were grown for at least five passages to separate tumor cells from tumor-associated fibroblasts. For signaling studies, cells were grown in serum-free conditions overnight, treated with 100 ng/mL recombinant murine leptin (R&D Systems) in serum-free media, and lysed for protein analysis after 20 minutes (peak induction of downstream signaling). Viability was assessed 72 hours after treatment with 100 ng/mL leptin using the CellTiter Glo (CTG) assay (Promega).

### Generation of leptin receptor knockout mPDAC cells

A single guide RNA (sgRNA) sequence (5’-TGAAAGCCACCAGACCTCGA) targeting exon 8 of the murine leptin receptor (LepR) was ligated into the BsmBI site in *lentiGuide-Puro* (Addgene 52963) with compatible annealed oligos to generate *lentiGuide-Puro-sgLepR*. Lentivirus for *lentiGuide-Puro-sgLepR* and *lentiCas9-blast* (Addgene 52962) were produced by co-transfection of 293T cells with lentiviral backbone, packaging vector (psPAX2), and envelope (VSV-G) using TransIT-LT1 (Mirus Bio). Supernatant was collected at 48 and 72 hours and applied to target cells with 8 μg/mL polybrene (EMD Millipore) for transduction. Transduced 7307 mPDAC cells were treated with 10 μg/mL blasticidin S (Life Technologies) and 2 μg/mL puromycin (Life Technologies) for 7 days. Cells were then sorted into single cells by limiting dilution into 96-well plates to isolate clones. Knockout clones were confirmed by genomic DNA extraction using QuickExtract DNA extraction solution (Epicentre), PCR amplification of the target locus (forward primer: 5’-GGTTCTCAGTGCACGCATTT-3’; reverse primer: 5’-ACAACGATTTTCCTGGCATCT-3’) with Q5 polymerase (NEB), and Sanger sequencing of PCR products by the Keck Biotechnology Resource Laboratory at Yale. KO1 clone harbored two frameshift mutant alleles: 32 bp deletion and an 18 bp deletion with a 2 bp insertion (effective 16 bp deletion). KO2 clone also harbored two independent frameshift mutant alleles: 1 bp deletion and a 13 bp deletion.

### Immunoblotting

Cells were lysed with ice-cold RIPA buffer (Thermo Fisher Scientific), supplemented with 0.5 μM EDTA and Halt protease and phosphatase inhibitors (Thermo Fisher Scientific), rotated at 4°C for 15-30 minutes to mix, and centrifuged at maximum speed for 15 minutes to collect whole cell lysates. Protein concentration was measured with the BCA protein assay (Pierce). 30 μg of total protein per sample was loaded into 4-12% NuPAGE Tris-Bis (Thermo Fisher Scientific) or 4-20% Tris-Glycine TGX (Bio-Rad) gradient gels and separated by SDS-PAGE. Proteins were transferred to nitrocellulose (for fluorescence detection) or PVDF (for chemiluminescent detection) membranes and blocked with Odyssey Blocking Buffer (LI-COR) or 5% milk, respectively. Primary antibodies used for immunoblotting are listed above. HSP90 and beta-tubulin were used as loading controls. Primary antibodies were detected with fluorescent DyLight-conjugated (Cell Signaling Technologies) or HRP-conjugated (BioRad) secondary antibodies for fluorescent (LI-COR Odyssey scanner) or chemiluminescent detection (Perkin Elmer ECL), respectively. Quantification of phosphorylated protein levels was performed using Image Studio Lite (LI-COR) and normalized to total protein levels.

### DNA and RNA isolation from mouse tumors

Mice were euthanized by cervical dislocation and pancreata were rapidly dissected, flash frozen in liquid nitrogen, and stored at −80C. Prior to DNA and RNA isolation, frozen tumors were mechanically ground in liquid nitrogen using a mortar and pestle. Genomic DNA and total RNA were extracted from ground tumor tissue using a Qiagen Allprep kit per manufacturer’s instructions. DNA concentration was measured using Picogreen (Invitrogen). RNA concentration, quality, and purity were determined using an Agilent Bioanalyzer.

### Mouse tumor exome sequencing and analysis

Exome capture on mouse tumor genomic DNA was performed by Macrogen Corp. using an Agilent Sureselect Mouse All Exon kit. 100-nt paired-end sequencing to a raw sequence depth of 150X per sample on a HiSeq 2500 (Illumina) was subsequently performed. Variant calls were made using an approach similar to the HaJaVa platform (McFadden et al., 2016). Reads were processed to remove adapter sequences (using exact matches to 10mer adapter start sequence: AGATCGGAAG) using the FASTX-toolkit (http://hannonlab.cshl.edu/fastx_toolkit). Surviving reads greater than or equal to 15nt in length were retained. Reads were aligned to the mouse mm9 genome assembly (UCSC) using BWA (v0.5.5). Duplicates were removed using Picard (v1.21) MarkDuplicates utility. Local realignment of reads around indels and base quality recalibration were performed using GATK (v1.0.5538). Variant calls were made with GATK UnifiedGenotyper using the mm9 reference sequence in regions targeted by the Agilent capture kit (version G7550_20110119_mm9 from Agilent). Calls with at least 14X read coverage and dual strand support were retained and further filtered using GATK VariantFiltration (arguments - clusterWindowSize 10 - cluster 3), GATK SelectVariants (arguments “SB < −0.1”, “QUAL >= 30.0”, “QD >= 5.0”, “HRun <= 5”), and Sanger Mouse Genomes Project Variant calls (Keane et al., 2011). Variants were annotated using Annovar (Wang et al., 2010) (version 2016Feb01) and mouse dbSNP build 142 (http://ftp.ncbi.nlm.nih.gov/snp/).

### Mouse tumor RNA sequencing (RNA-seq) and analysis

cDNA libraries from tumor total RNA were prepared by the MIT BioMicro Center using the Neoprep library preparation system (Illumina) with indexed adaptor sequences and polyA selection. Sequencing was performed on an Illumina NextSeq to obtain paired-end 75-nt reads. All reads that passed quality metrics were mapped to UCSC mm9 mouse genome build (http://genome.ucsc.edu/) using RSEM (v1.2.12) (http://deweylab.github.io/RSEM/). All RNA-seq analyses were conducted in R. High-resolution signature analyses between tumors from obese (*KCO* and *KCO* treated with AAV-GFP) and non-obese (*KC* and *KPC*) models were performed using a blind source separation methodology based on Independent Component Analysis (ICA), as previously described (Hyvarinen and Oja, 2000; Muzumdar et al., 2017).

Gene Set Enrichment Analyses (GSEA) (http://software.broadinstitute.org/gsea/) were carried out using standardized signature correlation scores (for ICA signatures) with default settings. Network representations of GSEA results were generated using EnrichmentMap (http://www.baderlab.org/Software/EnrichmentMap) for Cytoscape v3.3.0 (http://www.cytoscape.org) with p<0.005 and FDR<0.1. Each circle represents a gene set with circle size corresponding to gene set size and intensity corresponding to enrichment significance. Red is upregulated and blue is downregulated. Each line corresponds to minimum 50% mutual overlap with line thickness corresponding to degree of overlap. Cellular processes for gene set clusters were manually curated.

Tumor suppressor gene mutations in RNA-seq datasets were called using the GATK Best Practices workflow for SNP calling on RNA-seq data (software.broadinstitute.org/gatk/documentation/article.php?id=3891). Transcriptomic reads were trimmed to eliminate adapter sequences using an exact 10mer match to the start of the adapter sequence (AGATCGGAAG) using Cutadapt (v1.16) and subsequently mapped to the mouse mm9 (UCSC) genome assembly using the STAR RNA-seq aligner with default parameters. Picard v2.17.0 MarkDuplicates utility was used to mark duplicates in the aligned data. GATK (v3.8.0) toolkit was used to “SplitNTrim” and reassign mapping qualities as per GATK best practices, and indel realignment and base recalibration were performed as recommended. Variants were called against the mm9 reference sequence using the GATK HaplotypeCaller and filtered using GATK VariantFiltration with recommended parameters. Variant calls were annotated using Annovar (version 2016Feb01).

Genes with standardized signature correlation scores z > 3 (alternatively z < −3) were used as gene sets to score TCGA (https://tcga-data.nci.nih.gov/tcga/) Pancreatic Adenocarcinoma (PAAD) tumors (The Cancer Genome Atlas Research Network, 2017) using ssGSEA (Barbie et al., 2009). Tumors were stratified based on standardized signature scores (z-scores) derived using ssGSEA. Associated patients within top and bottom buckets (> or < 1 s.d.) were each assessed for over-representation of previously established molecular subtypes (squamous, immunogenic, progenitor, ADEX (Bailey et al., 2016)) using the hypergeometric test.

### Evaluation of cancer driver genes in human PDAC biospecimens

Acquisition of resected PDAC biospecimens including institutional review board approval and informed consent procedures were previously described (Qian et al., 2018). Pre-operative body-mass index (BMI) and molecular data on the main PDAC driver genes was available for 184 of 356 patients from this prior study. Of note, PDAC patients experience significant weight loss in the months preceding diagnosis, and weight at surgery may therefore not be fully reflective of weight in the prediagnostic period. In 125 patients with available weight data at 6 months before diagnosis, we compared weight at surgery with weight 6 months before diagnosis, and observed a strong correlation (Spearman’s rank correlation coefficient = 0.90, p<0.0001). This suggests that although weight may decrease in months preceding diagnosis, the rank order of 2 measurements is likely to be highly consistent. Targeted exome sequencing and immunohistochemistry (IHC) have been previously described (Qian et al., 2018). We used World Health Organization categories to classify patients as normal weight (<25 kg/m^2^), overweight (25-30 kg/m^2^), or obese (>30 kg/m^2^).

### Murine islet glucose stimulated insulin secretion (GSIS) studies

Islets from C57/B6 wild-type and *ob/ob* mice (n=4 mice per group) were isolated and perfused as previously described with a few minor changes, which are noted (Jesinkey et al., 2019). Isolated islets were recovered overnight and were perfused the following day in DMEM (D5030, Sigma) supplemented with 24mM NaHCO3, 10mM HEPES, 2.5mM glucose, 2mM glutamine and 0.2% BSA. The islets first equilibrated for 1hr at 2.5mM glucose on the perifusion instrument. After the stabilization period, islets were perfused with basal (2.5mM) glucose for 10 minutes followed by stimulatory glucose (16.7mM) for 45 minutes. After stimulation with glucose, the islets were exposed to basal glucose for 15 minutes followed by a final 30mM KCl step to ensure that the insulin secreting machinery distal to mitochondrial metabolism was intact in the two groups tested. During the perifusion, eluent was collected into a 96-well plate format and both secreted and total islet insulin concentrations were determined by a high range rodent insulin ELISA assay kit (ALPCO) and normalized to islet DNA using a PicoGreen dsDNA Quantitation Reagent Kit (Life technologies).

### Single cell RNA-sequencing of pancreatic islets

Murine pancreatic islets used for single cell analysis were isolated as described for the GSIS studies, however, were dispersed into single cells immediately after isolation using accutase (Gibco) and resuspended in 1 mL of PBS. Dead cells were removed using a MACS Dead Cell Removal Kit (Miltenyi Biotec) and cell concentration determined using a Countess II Automated Cell Counter (Thermo Scientific). Single cell library preparation was performed using a Chromium Single Cell 3’ Reagent Kit v3 (10X Genomics) and sequenced by Illumina HiSeq. Gene counts matrices were generated using CellRanger. Genes detected in fewer than 15 cells were removed and cells with library sizes greater than 15,000 UMI/cell were removed as potential doublets. We observed a bimodal distribution of mitochondrial gene detection and removed cells in the top 12.5% of library-size normalized mitochondrial expression as apoptotic cells. We then focused our analysis on single-hormone expressing islet cells as measured by expression of *Ins1/2*, *Gcg*, *Sst*, and *Ppy*, removing dual hormone-expressing cells and cells expressing markers of acinar, ductal, endothelial, or immune cells.

Cells were visualized using PHATE (Moon et al., 2019) with default parameters and gene expression was denoised using MAGIC (van Dijk et al., 2018) with default parameters. GSEA analyses were carried out comparing gene lists from scRNA-seq generated as described in the figure legend and applied to gene ontology sets (C5 biological processes and cellular components) in MSigDB (http://software.broadinstitute.org/gsea/) with default settings. To identify the association between gene-pairs, kNN-DREMI (van Dijk et al., 2018) was applied to selected genes and *Cck*. kNN-DREMI is an estimate of the mutual information which provides information about how much knowing the value of gene Y depends on the value of gene X even when the relationship between the two genes is non-linear. The DREMI scores between all three presented genes are in the top quartile of genes associated with *Cck*, with *Slc30a8* in the top 5%. Statistical testing for differential expression was performed using a Mann-Whitney U test as implemented in diffxpy (https://github.com/theislab/diffxpy).

### Evaluation of CCK expression in human biospecimens

IHC for detection of CCK expression in human PDAC samples was performed on 4 µm sections of formalin-fixed paraffin-embedded tissue blocks containing neoplastic and non-neoplastic pancreatic parenchyma that were obtained as part of specimens resected for pancreatic adenocarcinoma treatment, as previously described (Qian et al., 2018). Antigen retrieval was performed using citrate buffer (pH 6.0, Dako) in a 2100 Antigen Retriever system (Electron Microscopy Sciences). Sections were then incubated with dual endogenous peroxidase block (Dako) for 20 minutes at room temperature, protein block (Dako) for 10 minutes at room temperature, and anti-CCK8 primary antibody for 1 hour at 4°C. Chromogenic visualization was subsequently performed using the EnVision+ System-HRP (Dako) with a 3 minute diaminobenzidine (DAB) incubation followed by counterstaining with hematoxylin. For evaluation of CCK expression, the surrounding exocrine pancreas served as an internal negative control. If present within the section, neuroendocrine cells within the epithelium of the duodenum served as positive internal controls. The immunohistochemical assessment of CCK expression was conducted without knowledge of BMI or other patient data.

Human chronic pancreatitis (etiologies included pancreatic divisum (n=5), Sphincter of Oddi dysfunction (n=2), idiopathic (n=5)) specimens were obtained from 12 adults (ages 25-60, M:F 4:8) undergoing Total Pancreatectomy and Islet Auto-Transplant (TP-IAT) program at the University of Minnesota Medical Center. Pancreatic biopsies were obtained at the time of surgery and processed for routine histologic analyses. Informed consent was obtained from all patients, and the research protocol was reviewed and approved by the University of Minnesota Institutional Review Board. A total of 127 islets were evaluable, of which 4 harbored rare CCK-positive cells.

108 human islets of northern European descent were isolated at the Oxford Consortium for Islet Transplantation (OXCIT, Oxford, UK) as previously described (Cross et al., 2012). All studies were approved by the University of Oxford’s “Oxford Tropical Research Ethics Committee” (OxTREC Reference: 2-15), or the Oxfordshire Regional Ethics Committee B (REC reference: 09/H0605/2). All organ donors provided informed consent for use of pancreatic tissue in research. RNA was extracted, sequenced, and analyzed as previously described(van de Bunt et al., 2015). Reads per kilobase of CCK transcript per million mapped reads (RPKM) was log2 normalized for analysis. For a subset of donors, BMI was not available, and samples could not be used in analysis. Given limited sample size, low and high BMI groups were specified based on median United States BMI (28.2 kg/m^2^) defined by the most recent 2015-2016 National Health and Nutrition Examination Survey (NHANES) (https://www.cdc.gov/nchs/nhanes/index.htm). This also closely approximated median BMI in the donor cohort of 28 kg/m^2^

### Induction of murine pancreatitis

Acute pancreatitis was induced in adult C57/B6 mice (∼25 grams in weight) using cerulein (Sigma; six hourly 50 µg/kg intraperitoneally (IP) injections) or arginine (Sigma; three hourly 3 g/kg IP injections). Mice were euthanized and processed for histology at 12 and/or 48 hours after the first injection. Chronic pancreatitis was induced in adult C57/B6 mice with cerulein given six hourly 50 µg/kg IP injections three times a week for 2 weeks prior to analysis. Acute or chronic PBS-treated mice were used as controls. Two mice from each treatment group were analyzed.

### Data and code availability

The datasets generated during and/or analyzed during the current study are available in the NCBI Sequencing Read Archive (SRA) under accession number PRJNA544740 and the NCBI Gene Expression Omnibus (GEO) under accession numbers GSE131714 and GSE137236. Primary murine PDAC cell lines and constructs are available upon request. Computer code for RNA-seq signature analysis (ICA) and Python scripts for scRNA-seq analyses (PHATE and MAGIC) will be made available on GitHub or upon request. Other software tools (including version numbers) for exome and RNA-seq analyses are listed above or referenced.

### Statistical analyses

P-values for comparisons of two groups were determined by two-tailed student’s t-test (for normally distributed data) or Mann-Whitney U test (for non-parametric data) as noted in the figure legends using Prism (Graphpad). All replicates were included in these analyses and represent measurements from distinct samples (biologic replicates). The log-rank test was used for Kaplan-Meier survival analyses with Prism. Using SAS software (SAS Institute, Inc., Version 9.4), we performed logistic regression adjusted for age at surgery (continuous), gender (men, women), and study site to evaluate the association between BMI (normal, overweight, obese) and driver alterations in human PDAC. We calculated odds ratios (ORs) and 95% confidence intervals (CIs). We used the Kruskal-Wallis test to compare the number of driver alterations across BMI categories. A p-value of <0.05 was used to denote statistical significance. All error bars denote standard error of mean (s.e.m.) unless otherwise denoted.

## SUPPLEMENTARY INFORMATION

**Figure S1.**
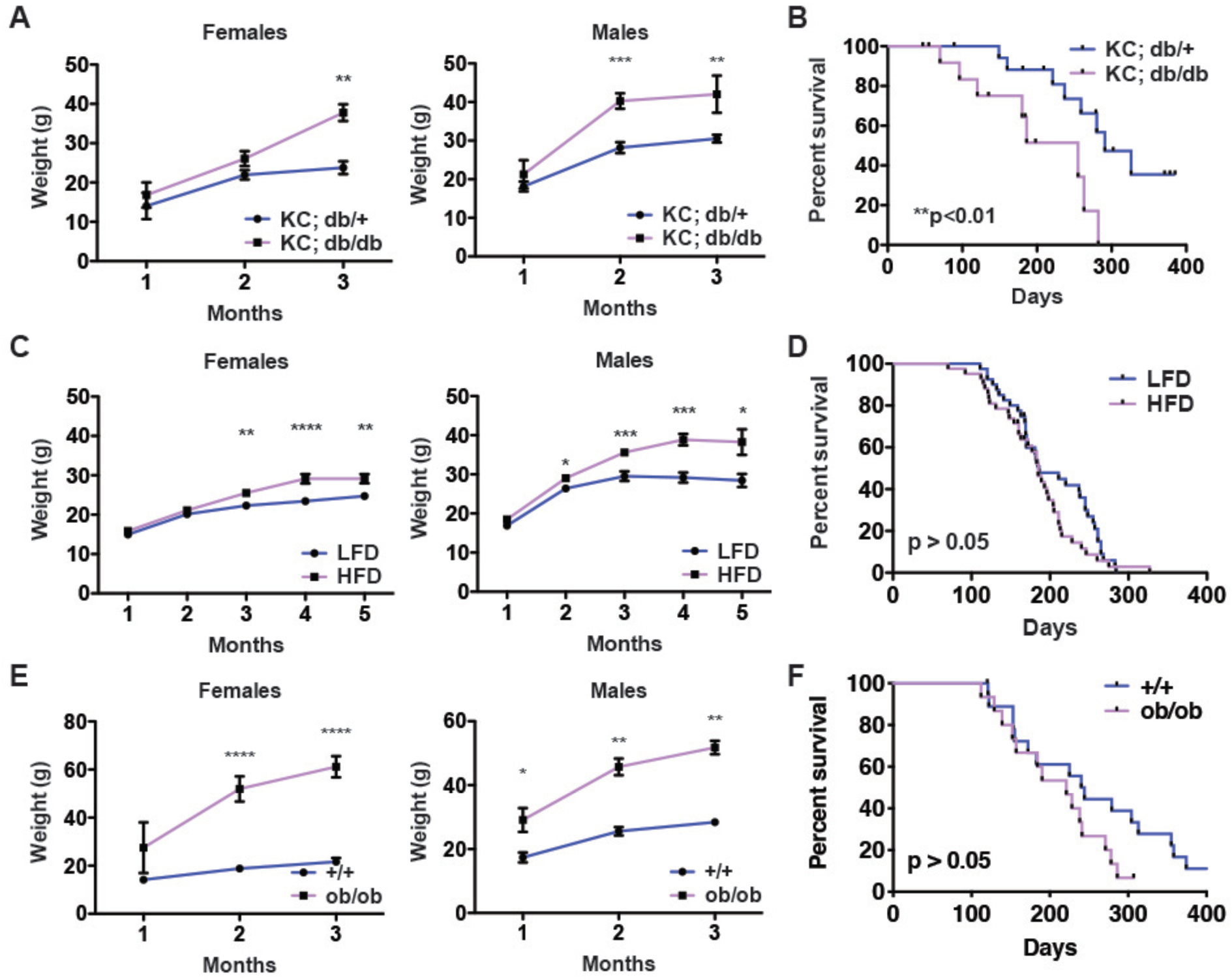
Obesity drives pancreatic but not lung tumorigenesis in mice, Related to Figure 1. A) Average body weight +/− s.e.m. (in grams) of female and male *KC* mice of varying *db* genotype over time (n=4-12 mice per group). B) Kaplan-Meier survival curves for non-obese (*db/+*, n=18) and obese (*db/db*, n=14) *KC* mice. C) Average body weight +/− s.e.m. (in grams) of female and male *Kras^LA2^* mice fed low-fat (LFD) or high-fat (HFD) diet (n=18-20 mice per group). D) Kaplan-Meier survival curves for LFD- (n=40) and HFD-fed (n=42) *Kras^LA2^* mice. E) Average body weight +/− s.e.m. (in grams) of female and male *Kras^LA2^* mice of varying *ob* genotype (n=3-9 mice per group). F) Kaplan-Meier survival curves for non-obese (*+/+*, n=19) and obese (*ob/ob*, n=15) *Kras^LA2^*mice. *p<0.05, **p<0.01, ***p<0.001, ****p<0.0001, two-tailed student’s t-test for all pairwise comparisons.

**Figure S2.**
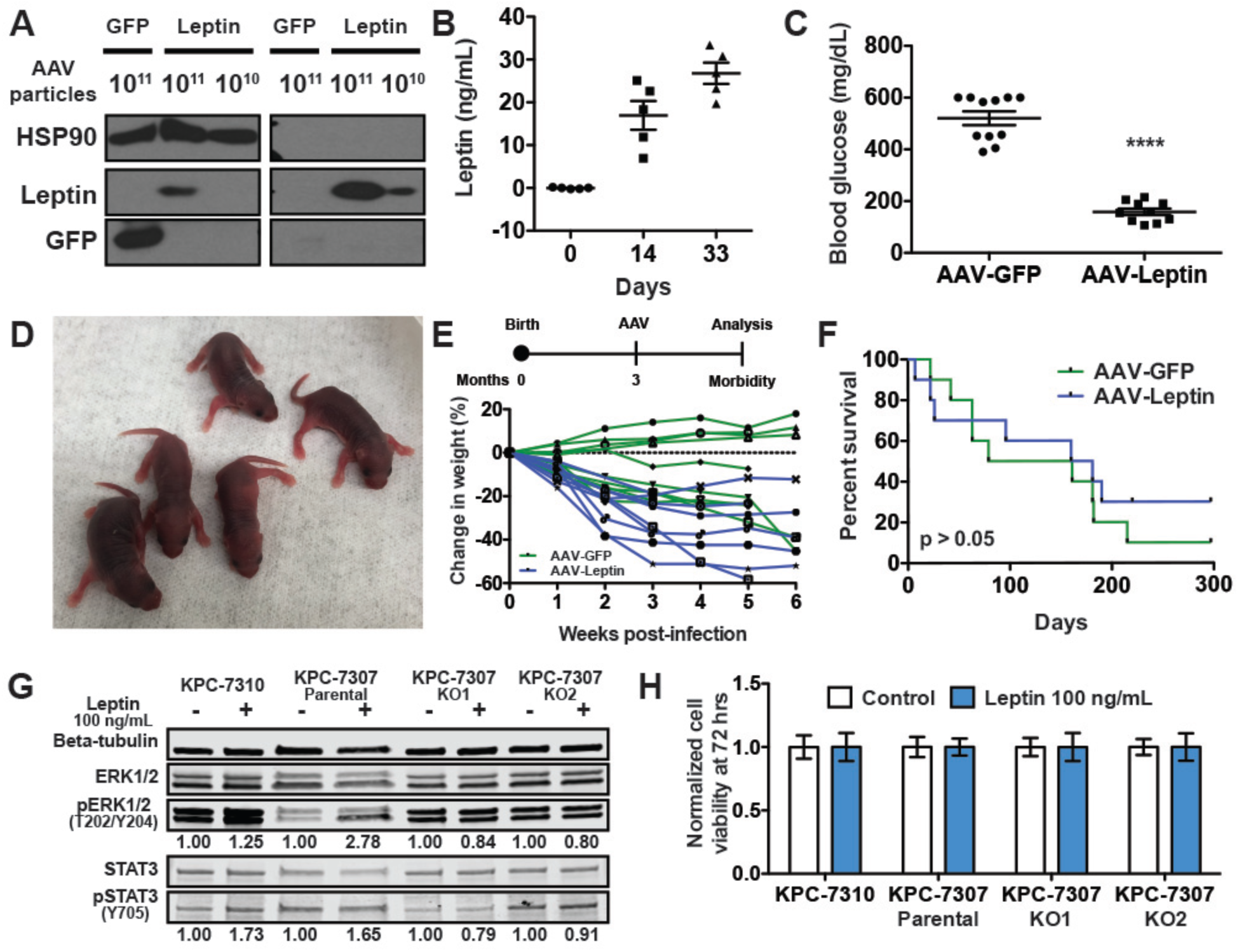
Adeno-associated virus-induced leptin expression in cells and mice, Related to Figure 2. A) Western blot showed dose-dependent leptin protein expression and secretion in 293T cells 48 hours after infection with AAV-Leptin or AAV-GFP control. HSP90 is cellular loading control. B) ELISA measurement of serum leptin concentrations (mean +/− s.e.m) at 0, 14, and 33 days following intramuscular injection of AAV-Leptin in adult *ob/ob* mice shows sustained expression. X-axis is time (in days) post-infection with AAV. C) AAV-Leptin induced a reduction in random blood glucose (mean +/− s.e.m.) 6 weeks post-infection of *KCO* mice. **** p<0.0001, two-tailed student’s t-test. Mice were infected at 6 weeks of age. D) AAV-Leptin restored fertility in *ob/ob* male and female mice. Shown are early postnatal progeny born from an AAV-Leptin-treated *ob/ob* female mated with an AAV-Leptin-treated *ob/ob* male. E) Schematic of AAV treatment of tumor-bearing mice. AAV is administered at 3 months of age once tumors have progressed. Percent change in body weight for each mouse (n=10 mice/group) following AAV administration is shown. F) Kaplan-Meier survival curves for mice in (E) shows no significant difference (p>0.05, log-rank test) between AAV treatments (n=10 mice/group). G) Recombinant murine leptin (100 ng/mL) induces MAPK (pERK1/2) and STAT3 (pSTAT3) signaling in murine *Kras*/*p53* mutant PDAC (mPDAC) cells (KPC-7307 and KPC-7310), which is abolished in leptin receptor knockout clones (KPC-7307 KO1 and KO2). Western blot shows protein levels at 20 minutes following treatment (peak induction). Quantification of pERK/ERK and pSTAT3/STAT3 levels (normalized to untreated) is shown. Beta-tubulin is additional loading control. Leptin dose was chosen to be within an order of magnitude of that observed in the serum of AAV-Leptin-treated mice in (b). H) Recombinant murine leptin (100 ng/mL) has no effect (p>0.05, two-tailed student’s t-test) on mPDAC cell viability (+/− s.d., normalized to untreated) 72 hours are treatment (n=5 replicates/condition).

**Figure S3.**
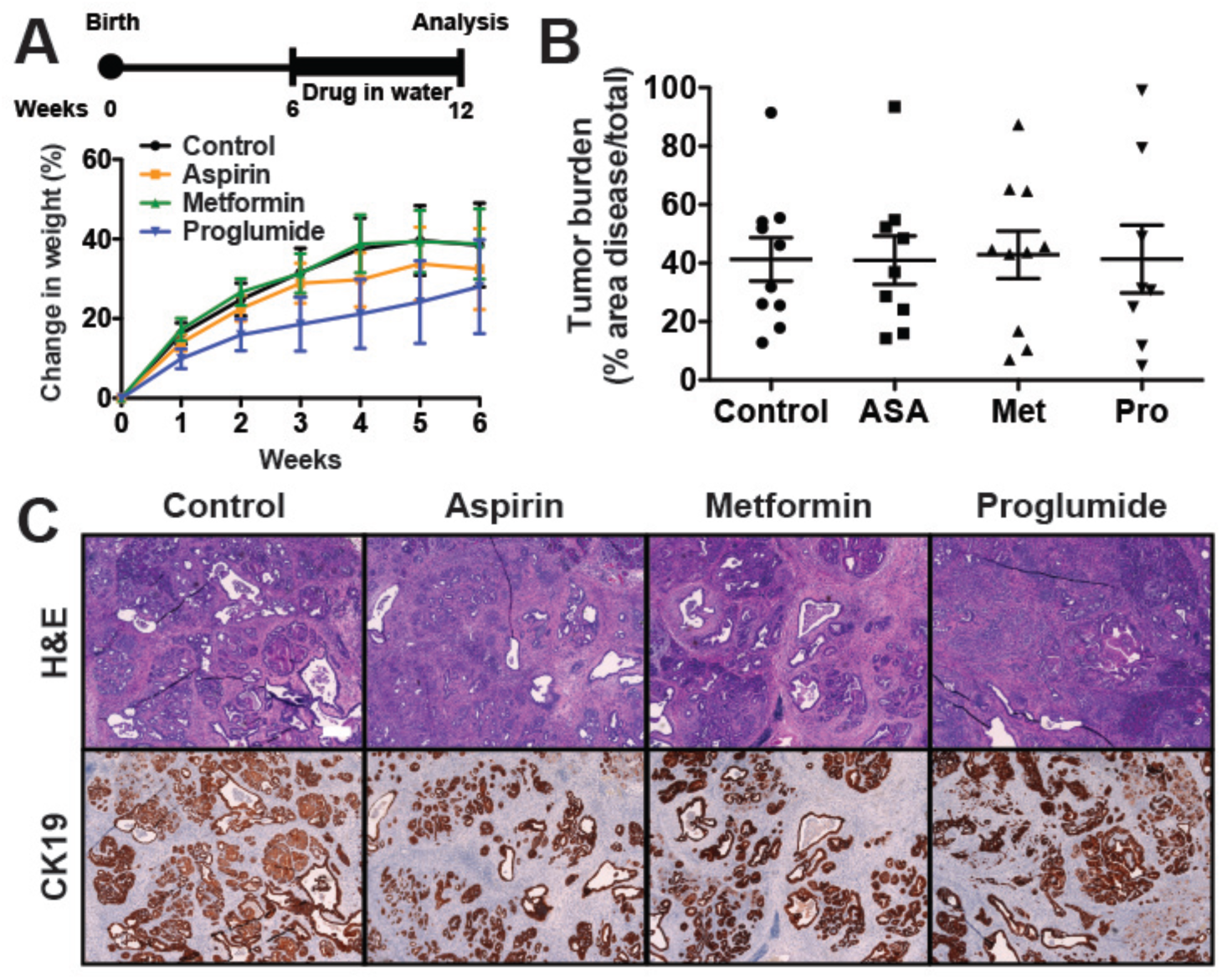
Treatment studies in *KCO* mice, Related to Figures 4 and 7. A) Schematic shows treatment strategy in which mice are placed on drinking water containing drug at 6 weeks and analyzed for tumor progression 6 weeks later. Percent change in body weight (mean +/− s.e.m.) are shown for each treatment (n=7-10 mice/group). The CCK receptor antagonist proglumide showed a modest slowing of weight gain. B) Quantification of tumor burden (average % cross-sectional area of disease/total +/− s.e.m.) is shown (n=7-10 mice/group) for each treatment (ASA = aspirin, Met = metformin, Pro = proglumide) and control drinking water. No significant differences were observed (p>0.05, two-tailed student’s t-test). C) Representative histologic sections of pancreata of mice at endpoint demonstrate comparable development of CK19-positive ductal tumors in all treatment groups in (b). Scale bar: 200 µm.

**Figure S4.**
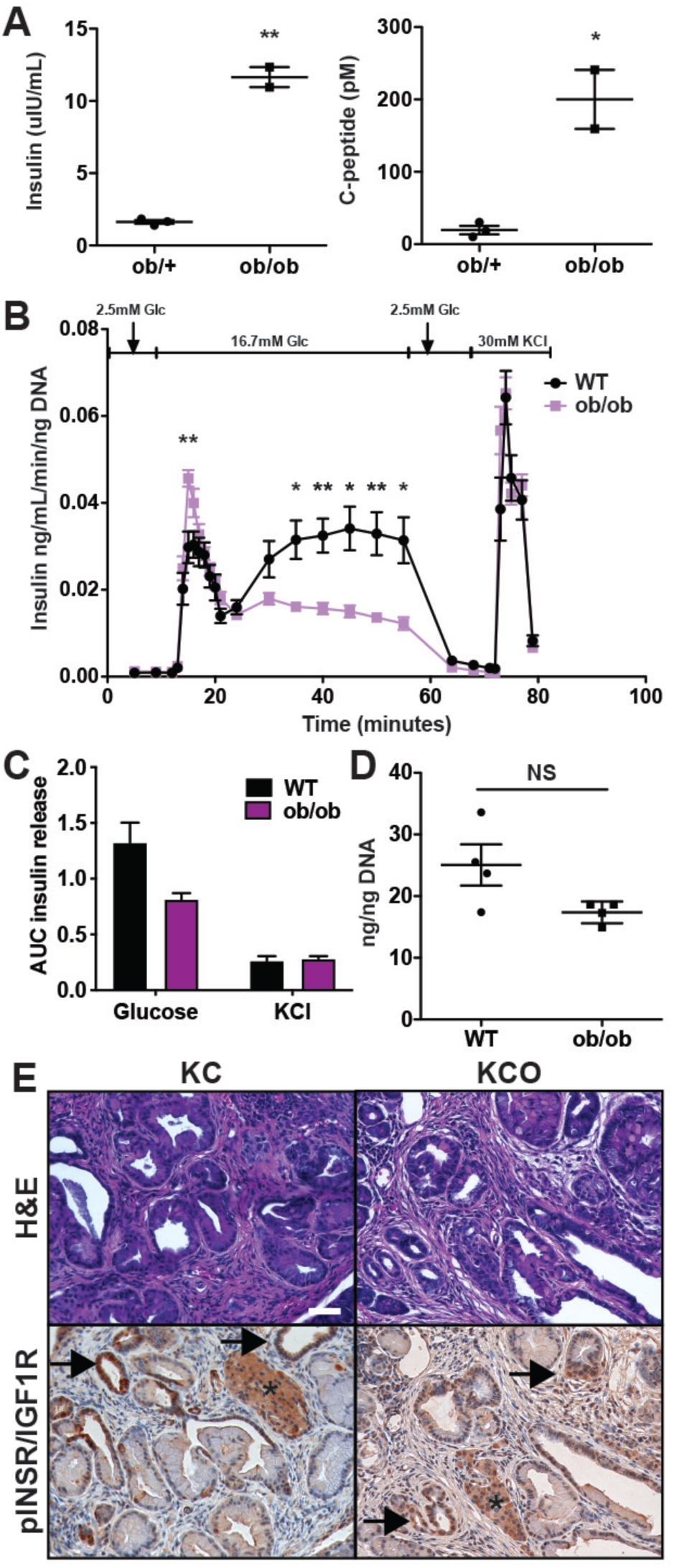
Insulin secretion and signaling alterations in obese mice, Related to Figure 5. A) Obese leptin-deficient (*ob/ob)* mice exhibited increased fasting serum insulin and C-peptide levels compared to non-obese controls (*ob/+*) at 10 weeks (n=2-3 mice/group). *p<0.05, **p<0.01, two-tailed student’s t-test. B) Glucose-stimulated insulin secretion (mean +/− s.e.m., n=4 replicates/group) from perifused islets derived from 12 week-old C57/B6 wild-type and *ob/ob* mice (n=4 mice/group). *p<0.05, **p<0.01, two-tailed student’s test for each timepoint. C) Area under the curve (AUC) (mean insulin ng/mL/ng DNA +/− 95% confidence intervals) shows reduced glucose-stimulated insulin secretion (combined 1^st^ and 2^nd^ phase) for *ob/ob* islets in (B). AUC for potassium chloride (KCl)-induced depolarization is not significantly different, suggesting no change in overall insulin stores. D) Total insulin content (mean +/− s.e.m., normalized to DNA content) in cultured islets in (B). E) IHC reveals activated insulin receptor/Igf1 receptor (phospho-InsR/Igf1R) in PanINs in non-obese (*KC*) and obese (*KCO*) mice, consistent with insulin-induced signaling. pInsR/Igf1R increases in higher-grade PanINs (arrows). Pancreatic islets were internal positive controls (asterisks). Scale bar: 50 µm.

**Figure S5.**
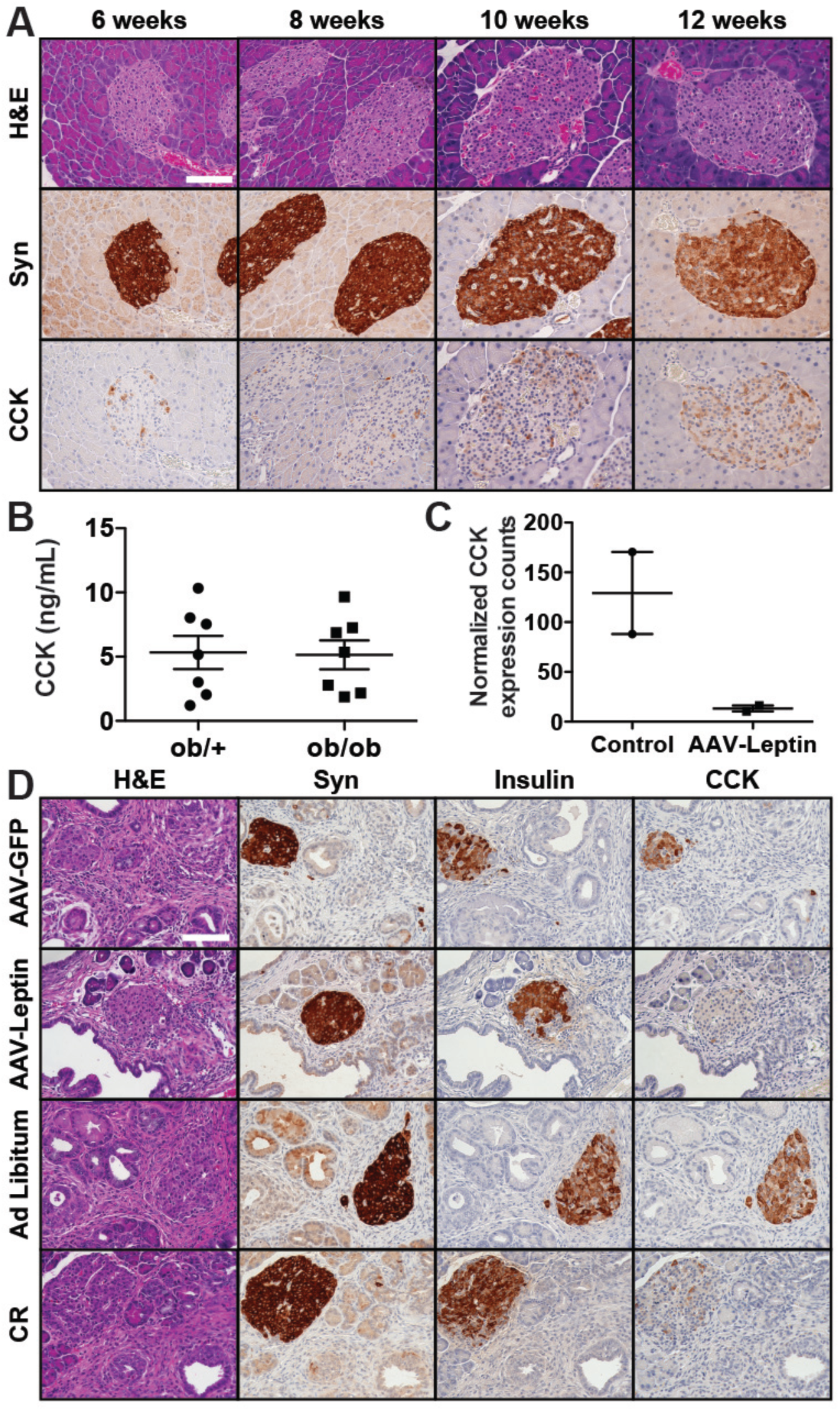
Obesity-associated islet CCK expression in mice, Related to Figures 6 and 7. A) Obese (*ob/ob*) mice exhibit progressive expression of CCK in pancreatic islets (denoted by Syn (synapotphysin) immunostaining). Scale bar: 100 µm. B) Non-fasting serum CCK levels (mean +/− s.e.m.) did not differ between 7-9 week-old obese *ob/ob* and non-obese *ob/+* mice (n=7 mice/group). C) AAV-leptin treatment of *ob/ob* mice reduced CCK expression by RNA-seq on bulk pancreata. Normalized CCK expression counts (mean +/− s.e.m). are shown (n=2 mice/group). D) Islet CCK expression is abolished following weight loss induced by AAV-Leptin or caloric restriction of *KCO* mice. Insulin denotes beta cells. Mice were analyzed at 12 weeks of age following intervention starting at 6 weeks of age. Scale bar: 100 µm.

**Figures S6.**
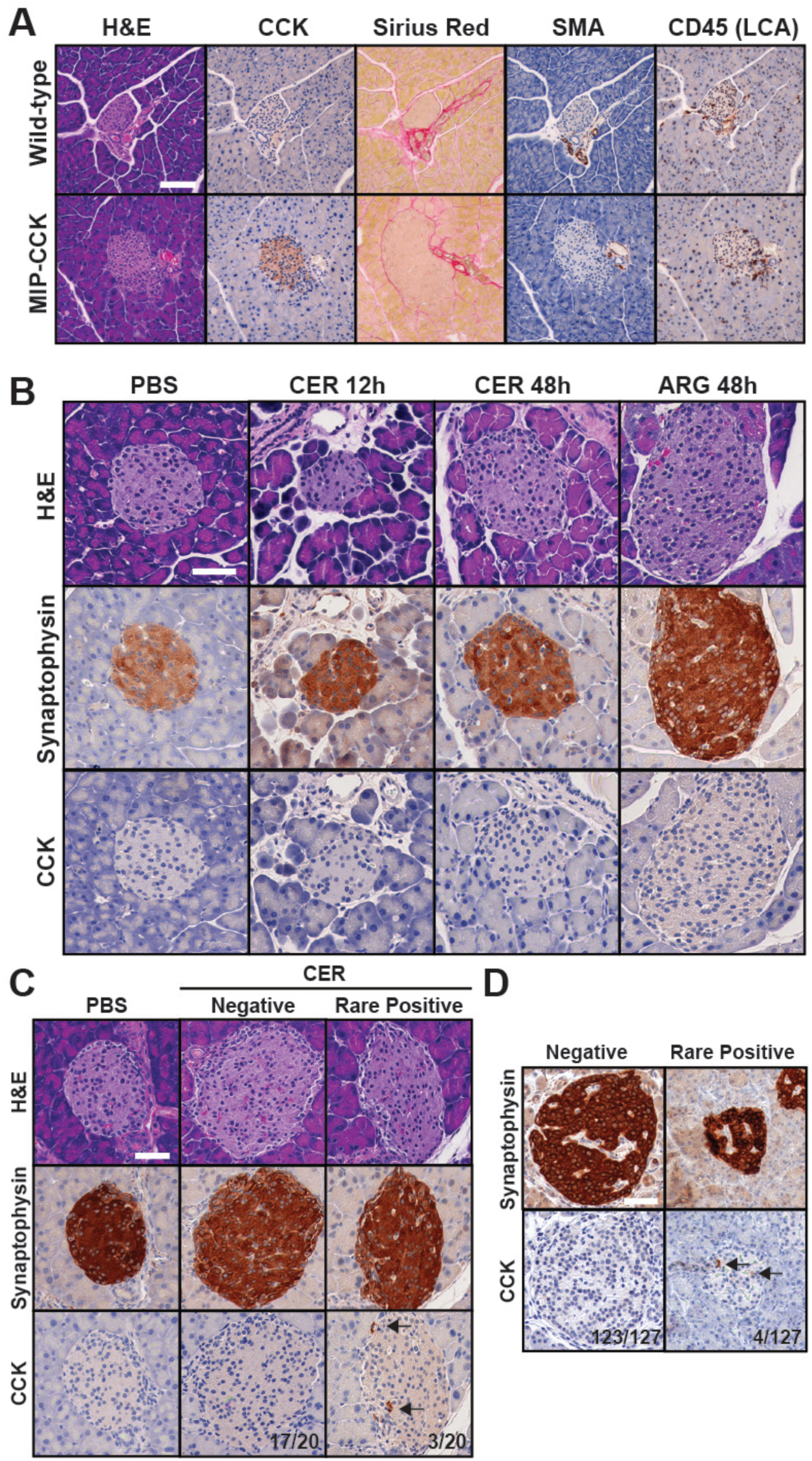
Disconnect between islet CCK expression and pancreatic inflammation, Related to Figure 7. A) *MIP-CCK* transgenic mice exhibited islet-specific CCK expression without evidence of pancreatic inflammation (CD45-positive leukocyte infiltration) or fibrosis (Sirius red collagen staining or presence of smooth muscle actin (SMA)-positive fibroblasts compared to wild-type controls. Scale bar: 100 µm. B) CCK is not significantly expressed in pancreatic islets in murine models of acute pancreatitis (CER = cerulein, ARG = arginine, PBS is control). Scale bar: 100 µm. C) CCK is rarely expressed (arrows) in pancreatic islets in a murine model of chronic pancreatitis (cerulein-treated for 2 weeks). Frequencies of islets without staining (negative) or rare CCK-positive cells (arrows) are shown. Scale bar: 100 µm. D) CCK is rarely expressed (arrows) in pancreatic islets obtained from patients with chronic pancreatitis. Frequencies of islets without staining (negative) or rare CCK-positive cells are shown. Scale bar: 100 µm.

**Table S1. Tumor suppressor gene (TSG) single nucleotide variants (SNVs) in *KCO* tumors, Related to Figure 3.**

Mutant allele fractions for each coding nucleotide (compared to mm9 reference database) of tumor suppressor genes (*Trp53*, *CDKN2A/p16*, and *SMAD4*) derived from whole exome sequencing and RNA-seq are shown for individual *KCO* tumors and averaged across all tumors Average base coverage across *KCO* tumors are as follows: *Trp53* (84X exome, 127X RNA), *CDKN2A/p16* (40X exome, 89X RNA), and *SMAD4* (103X exome, 86X RNA). Blank cells denote lack of base coverage in sequencing, which was rare and sporadic. A single *KPC* tumor (7075) was used as a positive control for SNV detection (resulting in a R172H codon substitution as noted). No tumor suppressor gene SNVs were detected in *KCO* tumors.

**Tables S2. Single nucleotide variants in murine tumors, Related to Figure 3.**

All single nucleotide variants (SNV) called in *KCO* (n=13) and *KPC* (n=1) tumors by whole exome sequencing are listed by tumor. Variant information includes location (chromosome, cytoband, and position in mm9 genome), genic location (exonic, intergenic, UTR), nucleotide change (var vs. ref), read counts for variant (tumor_reads2) and reference (tumor_reads1), and variant frequency (tumor_var_freq). Exon SNVs were further classified as synonymous or non-synonymous with reference to transcript (RefSeq), exon number (Exon_num), and coding sequence position (AA_change).

**Table S3. Compilation of non-synonymous single nucleotide variants in murine tumors, Related to Figure 3.**

Compiled non-synonymous single nucleotide variants (SNVs) identified on whole exome sequencing data derived from all sequenced *KCO* and *KPC* tumors from Table S2. A total of 531 SNVs were called from 14 tumors (∼37.7 SNVs per *KCO* mouse (n=13) and 41 SNVs in *KPC* mouse (n=1)). Induced *Kras*, *Lep* (*ob*), and *Trp53* mutations were observed as expected. Of consensus pan-cancer genetic drivers(Bailey et al., 2018), *Arid5b, Cacna1a, Eef2*, and *Kit* harbored recurrent non-synonymous SNVs. These SNVs were not observed in human cancer (COSMIC) to suggest pathogenicity.

**Table S4.** Characteristics of patients with available body mass index (BMI) and molecular alterations data, Related to Figure 3.

| **Patient characteristic** | **Mean (SD)** |
| --- | --- |
| Age at surgery, years | 65.5 (10.7) |
| BMI (kg/m<sup>2</sup>) | 26.7 (5.2) |
| Overall survival (months) | 23.1 (19.2) |
| Disease free survival (months) | 15.4 (17.2) |
| Tumor size (cm) | 3.6 (1.4) |
| **Sex** | **N (%)** |
| Men | 113 (61) |
| Women | 71 (39) |
| **Race** | **N (%)** |
| White | 144 (78) |
| Black | 24 (13) |
| Other/missing | 16 (9) |
| **Study site** | **N (%)** |
| DFCI/BWH<sup>1</sup> | 12 (7) |
| URMC<sup>2</sup> | 70 (38) |
| SCI<sup>3</sup> | 102 (55) |
| **Tumor location** | **N (%)** |
| Head/uncinate | 138 (75) |
| Body | 18 (10) |
| Tail | 25 (14) |
| Unknown | 3 (2) |
| **N stage** | **N (%)** |
| N0 | 55 (30) |
| N1 | 128 (70) |
| NX | 1 (1) |

<sup>1</sup>Dana-Farber Cancer Institute/Brigham and Women's Hospital
<sup>2</sup>University of Rochester Medical Center
<sup>3</sup>Stanford Cancer Institute

**Table S5.**
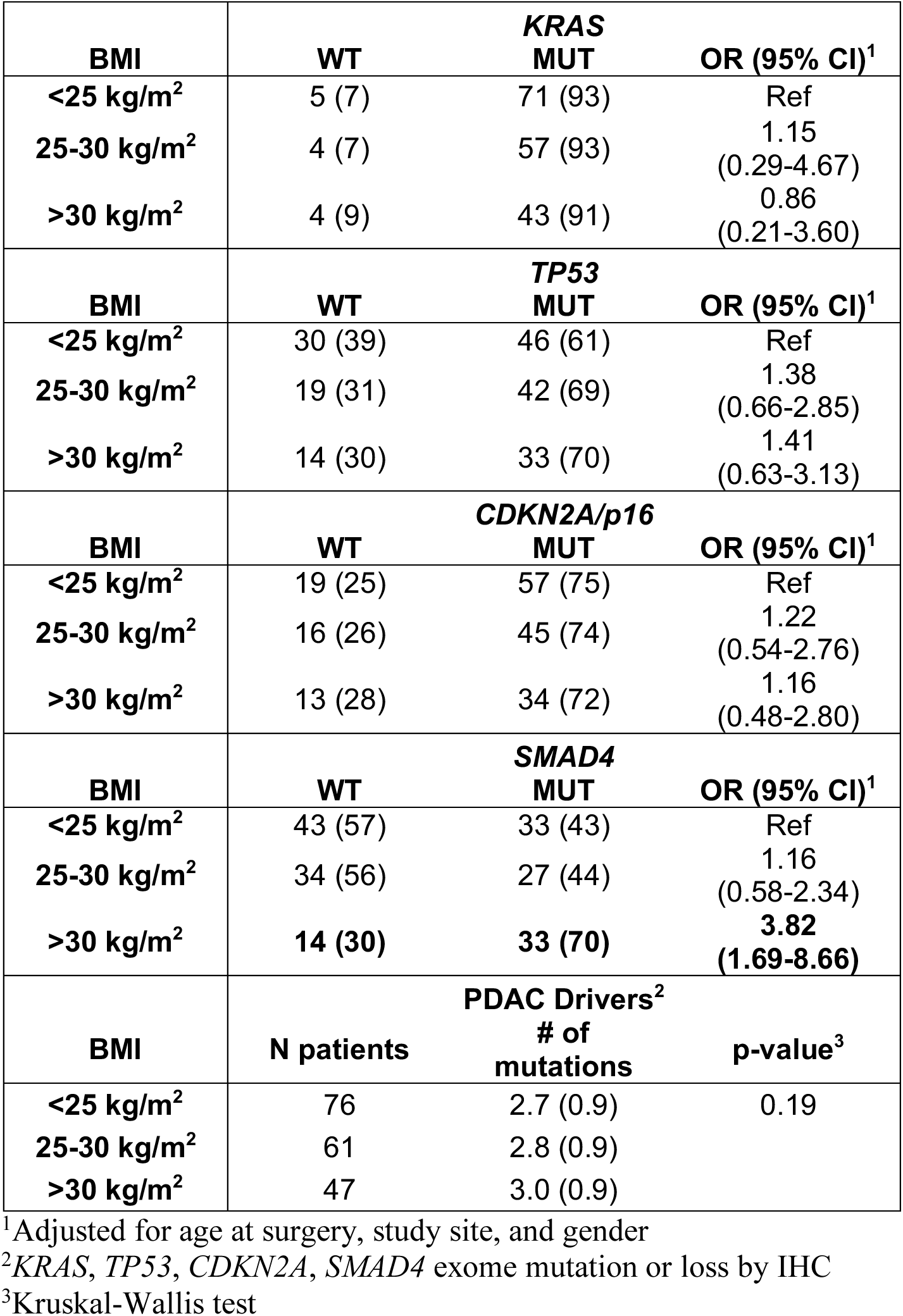
Association of BMI with PDAC driver gene alterations in human tumors, Related to Figure 3. Number (percentage of patients) at each BMI level are shown for each gene. For number of mutations, mean (standard deviation) is shown for each BMI level.

**Table S6. Normalized expression counts for RNA-seq of tumors, Related to Figures 4 and 5.**

Log (base 2) transformed normalized expression counts per gene in individual *KCO*, *KC*, and *KPC* tumors used in RNA-seq analyses. Mouse number, genotype, and AAV treatments are shown.

**Table S7. *KCO*-related gene signatures derived by independent component analysis (ICA), Related to Figures 4 and 5.**

Gene signatures that distinguish obese (*KCO*) and non-obese (*KC* and *KPC*) tumors by ICA with p<0.05 are shown. Per-gene expression correlation Z-scores with respect to the signature are listed (positive is correlated with *KCO* tumors) for each gene signature and ordered by log2 fold change comparing above groups.

**Table S8. Gene set enrichment analyses (GSEA) to determine *KCO*-related pathways, Related to Figures 4 and 5.**

GSEA results correlated with *KCO* tumors (UP) are shown comparing ICA-derived signatures (6, 12, and 13) in Table S7 with curated (C2) and hallmark (H) gene sets in MSigDB.

**Table S9. Beta cell comparisons from scRNA-seq data, Related to Figure 6.**

Composite gene expression differences of scRNA-seq data comparing *ob/ob* (OB) vs. wild-type (WT) beta cells or CCK-positive vs. CCK-negative OB beta cells. Shown are genes ordered by log2FC (fold change) including mean expression counts, p-values, and q-values, as measured using a Mann-Whitney U test implemented in diffxpy for each gene.

